# ANKRD49 promotes the metastasis of NCI-H1299 and NCI-H1703 cells via activating JNK-ATF2/c-Jun-MMP-2/9 axis

**DOI:** 10.1101/2023.03.22.533821

**Authors:** Jia Sun, Jin-rui Hu, Chao-feng Liu, Yuan Li, Wei Wang, Rong Fu, Min Guo, Hai-long Wang, Min Pang

## Abstract

Ankyrin repeat domain 49 (ANKRD49) has been found to highly expressed in multiple cancer including lung adenocarcinoma (LUAD) and lung squamous carcinoma (LUSC). However, the function of ANKRD49 in the pathogenesis of NSCLC still remains elusive. Previously, ANKRD49 has been demonstrated to promote the invasion and metastasis of A549 cells, a LUAD cell line, via activating the p38-ATF-2-MMP2/MMP9 pathways. Considering the heterogeneity of tumor cells, the function and mechanism of ANKRD49 in NSCLC need more NSCLC-originated cells to clarify. We found that ANKRD49 promoted the migration and invasion of NCI-H1299 and NCI-H1703 cells via enhancing the levels of MMP-2 and MMP-9. Furthermore, ANKRD49 elevated phosphorylation of JNK and then activated c-Jun and ATF2 which interact in nucleus to promote the binding of ATF2:c-Jun with the promoter MMP-2 or MMP-9. *In vivo* assay showed that ANKRD49 promoted lung metastasis of injected- NSCLC cells and the high metastatic rate was positively correlated with the high expression of ANKRD49, MMP-2, MMP-9, p-JNK, p-c-Jun and p-ATF2. In conclusion, the present study indicated that ANKRD49 accelerated the invasion and metastasis of NSCLC cells via JNK-mediated transcription activation of c-Jun and ATF2 which regulated the expression of MMP-2/MMP-9.

## 1. Introduction

Lung carcinoma is the most common cause of cancer-related deaths globally, with an estimated 1.6 million deaths annually(Bray et al., 2018). Non-small cell lung cancer (NSCLC) accounts for 85% of all cases(Keating, 2015), of which lung adenocarcinoma (LUAD) and lung squamous cell carcinoma (LUSC) are the most common subtypes(Huang et al., 2016). Notwithstanding unprecedented advances in the remedy of NSCLC, such as targeted therapy and immunotherapy, have been made in recent years, the long-term prognosis of NSCLC patients remains poor, with a 5-year survival rate of less than 15%(MacDonagh et al., 2018; Rosell and Karachaliou, 2013). Therefore, it is essential to clarify the molecular pathogenesis of NSCLC and identify more precise prognostic markers and therapeutic targets.

The ankyrin repeat domain consists of 30∼34 amino acid residues and mediates protein-protein interactions. ANKRD49 (ankyrin repeat domain 49) contains four ankyrin motifs. ANKRD49 is highly expressed in the testes and is involved in spermatogenesis(Wang et al., 2015; Zhou et al., 2019). ANKRD49 also participates in the progression of malignant gliomas and gastric cancer in humans(Hao et al., 2017; Liu et al., 2018). Previously, we explored the expression pattern of ANKRD49 protein in NSCLC (80 cases of LUAD and 80 cases of LUSC) and found that the levels of ANKRD49 in cancer tissues were higher than those in tumor-adjacent normal tissues, its expression correlated with the TNM (tumor-node-metastasis) stage, distal metastasis, lymph node metastasis and differentiation. Moreover, patients with higher ANKRD49 showed lower OS (overall survival) rate and higher ANKRD49 expression in lung tissues may serve as an independent prognostic marker for NSCLC patients(Li et al., 2022; Liu et al., 2022). Howbeit, the function and underlying mechanisms of ANKRD49 in NSCLC has not yet been fully elucidated.

Although, ANKRD49 has been demonstrated to promote the invasion and metastasis of A549 cells, a lung adenocarcinoma cell line, via activating the p38/ATF-2 signaling pathway and then elevating the levels of MMP2/MMP9 in our previous study(Liu et al., 2022). Considering the heterogeneity of tumor cells, the function and mechanism of ANKRD49 in NSCLC need more NSCLC-originated cells to clarify. Therefore, in the present study, another LUAD cell line (NCI-H1299) and a LUSC cell line (NCI-H1703) were selected and our findings illustrated that ANKRD49 accelerated the migration and invasion of NCI- H1299 and NCI-H1703 cells via the JNK-ATF2/c-Jun-MMP-2/MMP-9 pathway, unlike its function in A549 cells.

## 2. Results

### 2.1 ANKRD49 is highly expressed in NSCLC

To investigate the expression pattern of ANKRD49 in fresh tissues from NSCLC, nine fresh tumor tissues and adjacent normal tissues (6 cases of LUAD and 3 cases of LUSC) were collected and the ANKRD49 mRNA was measured using RT-qPCR. The data showed that the expression of ANKRD49 mRNA in tumor tissues was significantly higher than that in adjacent normal tissues (Fig. 1A). Next, the expression of ANKRD49 mRNA in human bronchial epithelial cell line (HBEC), as well as in five LUAD lines (H1299, A549, H446, H1460, Calu-3) and two LUSC lines (H1703, sk-MES-1) was assessed by RT-qPCR. The results demonstrated that the expression of ANKRD49 was obviously higher in H1299, A549, H446, H1703, and sk-MES-1 cells than in the human bronchial epithelial cell line (Fig. 1B). Accordingly, H1299 cell line was selected for ANKRD49 overexpression assay, while H1703 cell line was opted for ANKRD49 knockdown analysis.

**Fig. 1.**
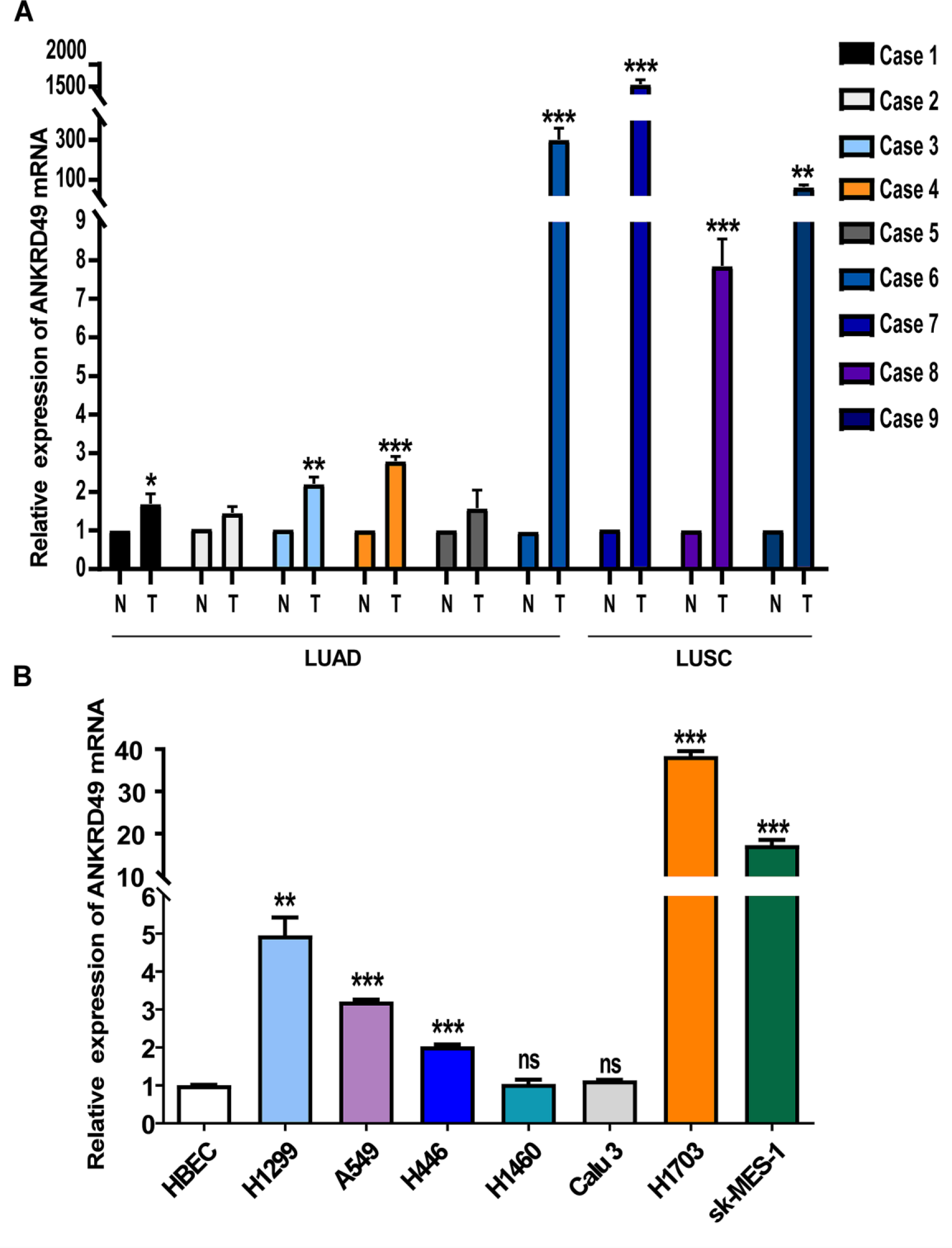
ANKRD49 is upregulated in NSCLC. (A) The mRNA levels of ANKRD49 in nine fresh NSCLC tissues and corresponding adjacent normal tissues were analyzed by RT-qPCR (N: normal tissues; T: tumorous tissues). (B) The mRNA levels of ANKRD49 were assessed by RT-qPCR in the human bronchial epithelial cell line (HBEC) and seven NSCLC cell lines. Data are expressed as means ± standard deviation. ns, not significant, \**P*<0.05, \*\**P*<0.01, \*\*\**P*<0.001 vs corresponding adjacent normal lung tissues or HBEC cells.

### 2.2 ANKRD49 promotes the migration and invasion of H1299 cells

To investigate the function of ANKRD49 in LUAD cells, we established stable ANKRD49-OE or ANKRD49-sh H1299 cells, as well as their respective control cells LV5 and LV3. RT-qPCR and Western blot assays showed that ANKRD49-OE and ANKRD49-sh H1299 cells were constructed (Fig. 2A, B). CCK-8 and colony formation assays revealed that ANKRD49 had no effect on H1299 cell proliferation (Supplementary Fig. S1). Wound healing, transwell migration, and invasion assays illustrated that ANKRD49-OE markedly enhanced the migration and invasion of H1299 cells compared to the LV5 group, and the opposite results were observed in the ANKRD49-sh cells compared to the LV3 group (Fig. 2C-F). Taken together, these data demonstrated that ANKRD49 accelerated the migration and invasion of H1299 cells.

**Fig. 2.**
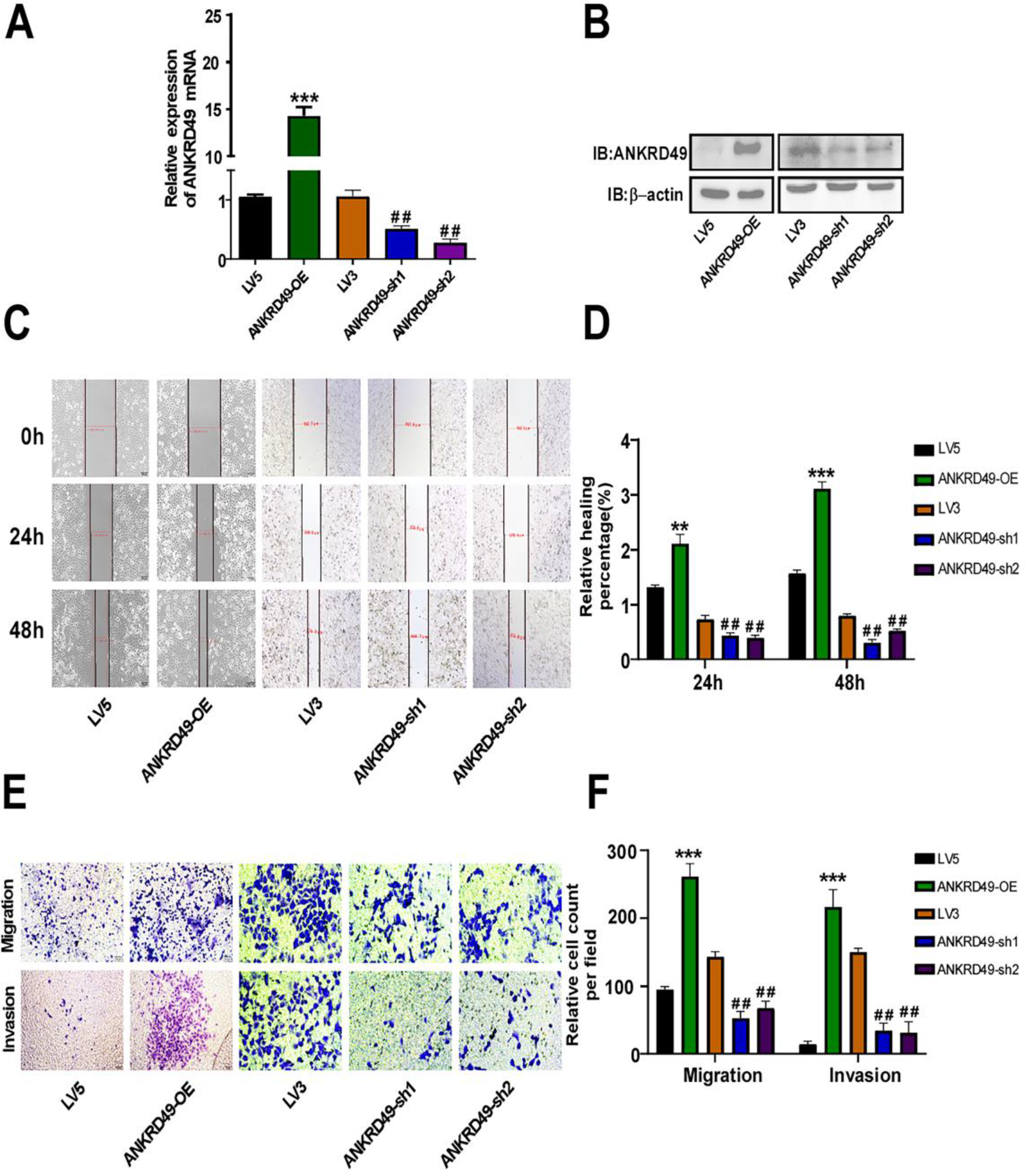
ANKRD49 potentiates migration and invasion of H1299 cells. (A, B) Identification of ANKRD49-OE or ANKRD49-sh H1299 cells was validated by RT-qPCR and Western blot. (C, D) Wound healing assay was used to measure migration of ANKRD49-OE and ANKRD49-sh H1299 cells, representative images were taken at a magnification of 40×, at 0, 24 and 48 h. (E, F) Transwell migration and invasion assays were performed to detect migration and invasion of ANKRD49-OE and ANKRD49-sh H1299 cells. Representative images were taken at 200× magnification. Statistical analysis from five random fields was conducted. All experiments were repeated independently three times. Data are expressed as means ± standard deviation. \**P*<0.05, \*\**P*<0.01, \*\*\**P*<0.001 vs LV5 group, ^#^*P*<0.05, ^##^*P*<0.01 vs LV3 group.

### 2.3 ANKRD49 upregulates MMP-2/MMP-9 expression by activating the MAPK pathway in H1299 cells

The high expression of MMP-2 and MMP-9 in cancerous tissues has been correlated with tumor cell matrix degradation, invasion, and migration(Feng et al., 2019; Wang et al., 2018; Xu et al., 2019). To uncover the molecular basis of ANKRD49 on invasion and migration, the expression of MMP-2 and MMP-9 was tested when ANKRD49 was overexpressed or downregulated by RT-qPCR and Western blot assays. The results revealed that MMP-2 and MMP-9 levels were elevated in ANKRD49-OE and attenuated in ANKRD49-sh H1299 cells compared with the LV5 and LV3 groups, respectively (Fig. 3A, B). Next, the gelatinase activities of MMP-2 and MMP-9 were markedly boosted in ANKRD49-OE H1299 cells and declined in ANKRD49-sh H1299 cells, compared with those in the LV5 and LV3 groups, respectively (Fig. 3C, D). In order to confirm whether the effect of ANKRD49 was dependent on MMP- 2 and MMP-9, ANKRD49-OE H1299 cells were pretreated with 10 µM ilomastat (MMP inhibitor) for 1 h. and the wound healing assay displayed that inhibition of MMP-2/MMP-9 attenuated the migration and invasion of H1299 cells (Fig. 3E, F).

**Fig. 3.**
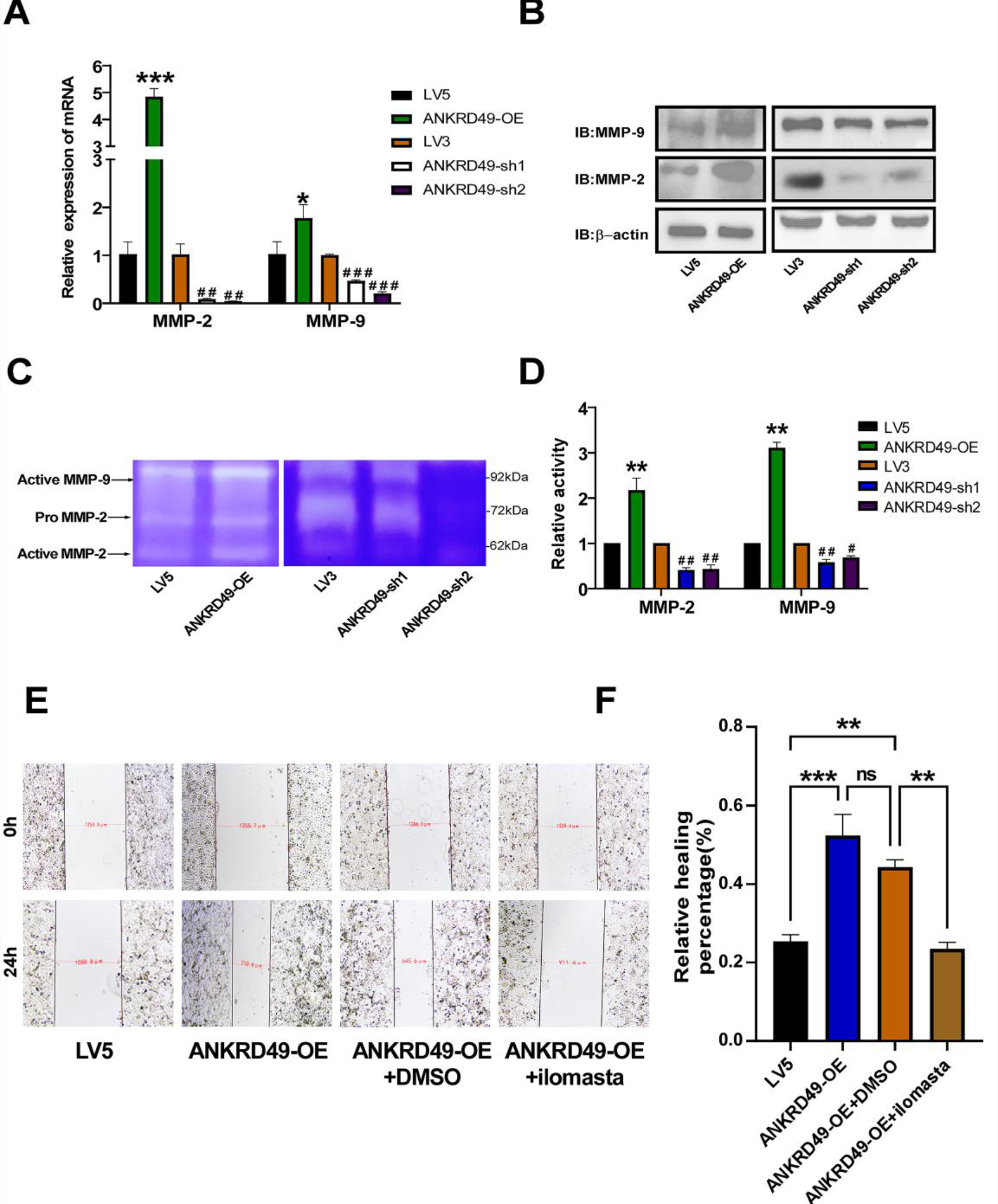
ANKRD49 upregulates MMP-2/MMP-9 expression in H1299 cells. (A, B) The mRNA and protein levels of MMP-2 and MMP-9 in ANKRD49-OE and ANKRD49-sh H1299 cells were detected using RT-qPCR and Western blot assays. (C, D) The activity of MMP-2 and MMP-9 in ANKRD49-OE and ANKRD49-sh H1299 cells was detected by gelatin zymography. (E, F) A wound healing assay was conducted to assess the effect of MMPs inhibitor (ilomastat) on the migration of ANKRD49-OE H1299 cells; representative images were taken at 40× magnification at 0 and 24 h. Data are expressed as means ± standard deviation. \**P*<0.05, \*\**P*<0.01, \*\*\**P*<0.001 vs LV5 group, ^#^*P*<0.05, ^##^*P*<0.01 vs LV3 group.

It is well established that mitogen-activated protein kinases (MAPKs), including p38 MAPK, ERK, and JNK, are involved in tumor metastasis and invasion(Li et al., 2020a; Wei et al., 2017; Zhou et al., 2020). As illustrated in Fig. 4A, the levels of p-p38 and p-JNK were obviously augmented in ANKRD49- OE and decreased in ANKRD49-sh H1299 cells compared with the LV5 and LV3 groups, respectively, while the levels of p-ERK were not altered. Next, p38 MAPK inhibitor (10µM SB203580), JNK inhibitor (10µM SP600125), or DMSO (10 µL) was used to pretreat ANKRD49-OE H1299 cells for 1 h. We found that SP600125 significantly decreased MMP-2/MMP-9 levels compared to DMSO-treated cells, whereas SB203580 had no effect on MMP-2/MMP-9 expression (Fig. 4B, C). Besides, the wound healing assay also exhibited that inhibition of JNK abated the migration and invasion of ANKRD49-OE H1299 cells, whereas inhibition of p38 had no effect on migration and invasion of ANKRD49-OE H1299 cells (Fig. 4D, E). To explore the specific role of JNK on MMP-2/MMP-9 expression, p-ATF2 and p-c-Jun, which are regulatory targets of JNK(Min et al., 2008), were analyzed by Western blot. As shown in Fig. 4F, the levels of p-ATF2 and p-c-Jun were both boosted in ANKRD49-OE and weakened in ANKRD49-sh H1299 cells compared with the LV5 and LV3 groups, respectively. Since ATF2 also has been reported to be regulated by p38 MAPK(Li et al., 2019), in order to clarify whether p38 MAPK was involved in the regulation of ATF2 in ANKRD49-OE settings, SP600125 or SB203580 was used and p-ATF2 was analyzed. The results illustrated that SP600125 decreased the level of p-ATF2, but SB203580 did not (Fig. 4G, H), indicating that p38 MAPK didn’t participate in the regulation of ATF2 in ANKRD49-OE H1299 cells. Overall, these data demonstrate that ANKRD49 promotes the invasion and migration of H1299 cells by enhancing MMP-2/MMP-9 expression mediated by the JNK but not p38 MAPK pathway.

**Fig. 4.**
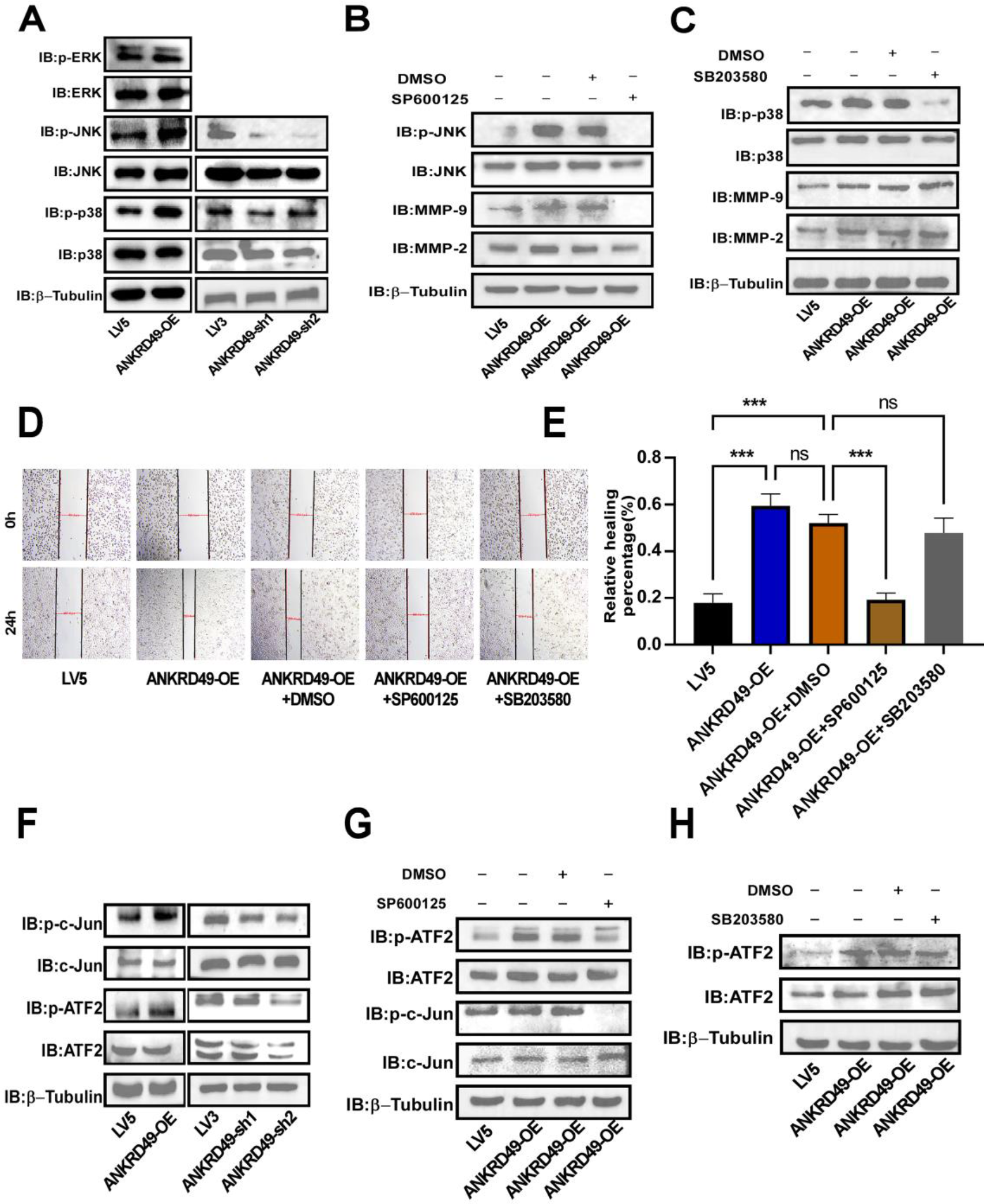
ANKRD49 activates JNK pathway to regulate the expression of MMP-2/MMP-9 in H1299 cells. (A) The MAPK protein levels in ANKRD49-OE and ANKRD49-sh H1299 cells were assessed by Western blot. (B, C) The effects of JNK inhibitor or p38 MAPK inhibitor on the levels of MMP-2/MMP- 9 in ANKRD49-OE H1299 cells were tested by Western blot. β-Tubulin served as an internal control. (D, E) A wound healing assay was conducted to assess the effect of JNK inhibitor (SP600125) or p38 inhibitor (SB203580) on the migration of ANKRD49-OE H1299 cells; representative images were taken at 40× magnification at 0 and 24 h. (F) The levels of p-ATF2 and p-c-Jun in ANKRD49-OE and ANKRD49-sh H1299 cells were measured by Western blot. (G, H) The levels of p-ATF2 and p-c-Jun in ANKRD49-OE H1299 cells treated with SP600125, SB203580 or DMSO were detected by Western blot. Data are expressed as means ± standard deviation. \**P*<0.05, \*\**P*<0.01, \*\*\**P*<0.001 vs LV5 group.

### 2.4 ANKRD49 activates ATF2/c-Jun transcription factor as a heterodimer to regulate MMP-2/MMP-9 expression in H1299 cells

ATF2 and c-Jun belong to the AP-1 family(Meng and Xia, 2011), translocate into the nucleus, and regulate the expression of target genes, including MMPs(Liu et al., 2020; Oh et al., 2020). Accordingly, the nuclear distribution of p-ATF2 and p-c-Jun was measured. The results showed that the nuclear levels of ATF2 and c-Jun as well as their phosphorylation were significantly enhanced in ANKRD49-OE H1299 cells compared to those in the LV5 group (Fig.5A, B). It is widely acknowledged that ATF2 contains a nuclear export signal (NES) in its leucine zipper region and two nuclear localization signals (NLS) in its basic region, leading to continuous shuttling between the nucleus and the cytoplasm(Lau and Ronai, 2012). It is essential for ATF2’s transcriptional activation that ATF2 dimerizes with c-Jun in the nucleus which prevents the export of ATF2(Gozdecka and Breitwieser, 2012). Co-immunoprecipitation assay revealed that ANKRD49 promoted the interaction between p-ATF2 and p-c-Jun in the nucleus (Fig. 5C). In addition, immunofluorescence assay also showed that ANKRD49 promoted the nuclear co- localization of p-ATF2 and p-c-Jun (Fig. 5D). Furthermore, chromatin immunoprecipitation assay was conducted to analyze the binding ability of ATF2 or c-Jun to the MMP-2 or MMP-9 promoter region. Our results illustrated that ANKRD49 enhanced the binding of ATF2 or c-Jun to the MMP-2 or MMP-9 promoter region (Fig.5E-H). Taken together, these findings suggest that ANKRD49 activates ATF2/c-Jun transcription factor as a heterodimer to regulate MMP-2/MMP-9 expression in H1299 cells.

**Fig. 5.**
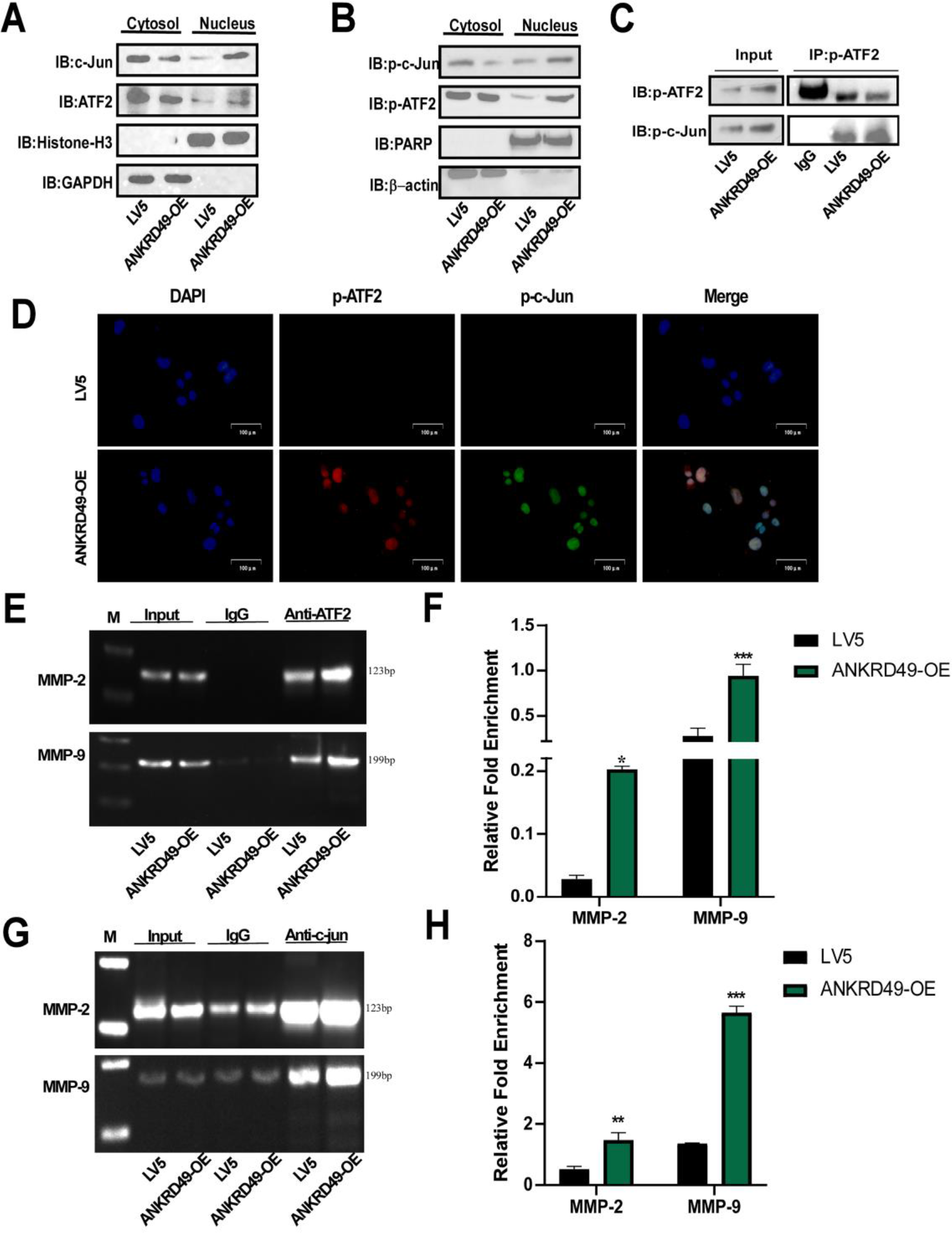
ANKRD49 activates c-Jun/ATF2 transcription factor to regulate the expression of MMP- 2/MMP-9 in H1299 cells. (A) The levels of ATF2 and c-Jun in the nuclear fraction of ANKRD49-OE or LV5 H1299 cells were measured by Western blot. (B) The levels of p-ATF2 and p-c-Jun in the nuclear fraction of ANKRD49- OE or LV5 H1299 cells were measured by Western blot. (C) Co-immunoprecipitation (Co-IP) analysis was performed to assess the interaction between p-ATF2 and p-c-Jun in the nucleus of ANKRD49-OE or vector H1299 cells. (D) Representative immunofluorescence images of ANKRD49 induced nuclear co-location of p-ATF2 and p-c-Jun. Scale bar:100 μm. (E-H) Chromatin immunoprecipitating (CHIP) assay was carried out to analyze the binding of ATF2 or c-Jun with the promoter of MMP-2 or MMP-9. β-Tubulin, and PARP or Histone-H3 served as the internal control for cytosol and nuclear fractions, respectively. Data are expressed as means ± standard deviation. \**P*<0.05, \*\**P*<0.01, \*\*\**P*<0.001 vs LV5 group.

### 2.5 ANKRD49 promotes migration and invasion of H1703 cells via MMP-2/MMP-9 mediated by JNK-ATF2/c-Jun pathway

Furthermore, the function of ANKRD49 in LUSC was investigated by establishing stable ANKRD49 knockdown (ANKRD49-sh) H1703 cells and a corresponding control LV3. RT-qPCR and Western blot assays demonstrated that ANKRD49-sh H1703 cells were successfully prepared (Fig. 6A, B). CCK-8 and colony formation assays revealed that ANKRD49 had no effect on H1703 cells’ proliferation (Supplementary Fig. S2). Wound healing, transwell migration, and invasion tests exhibited that ANKRD49-sh dramatically attenuated the migration and invasion of H1703 cells compared to the LV3 group (Fig. 6C-F). Additionally, we found that downregulation of ANKRD49 diminished both the expression and activity of MMP-2 and MMP-9 (Fig. 7A-D). Similarly, downregulation of ANKRD49 reduced the levels of p-JNK, p-p38, p-ATF2, and p-c-Jun in H1703 cells (Fig. 7E, F). To further confirm whether the effect of ANKRD49 on the MMP-2/MMP-9 levels in H1703 cells was dependent on JNK pathway just like it’s function in H1299cells, Anisomycin (20µM), an activator of JNK and p38 MAPK(Wang et al., 2022) was used to pretreat ANKRD49-sh H1703 cells for 1 h, followed by administration with SB203580, SP600125 or DMSO for another 24 h. We found that Anisomycin significantly augmented MMP-2/MMP-9 levels in SB203580-treated cells, whereas Anisomycin had no effect on MMP-2/MMP-9 expression in SP600125-treated cells (Fig. 7G, H). These data suggest that ANKRD49 promotes the invasion and migration of H1703 cells by enhancing MMP-2/MMP-9 expression mediated by the JNK rather than p38 MAPK pathway.

**Fig. 6.**
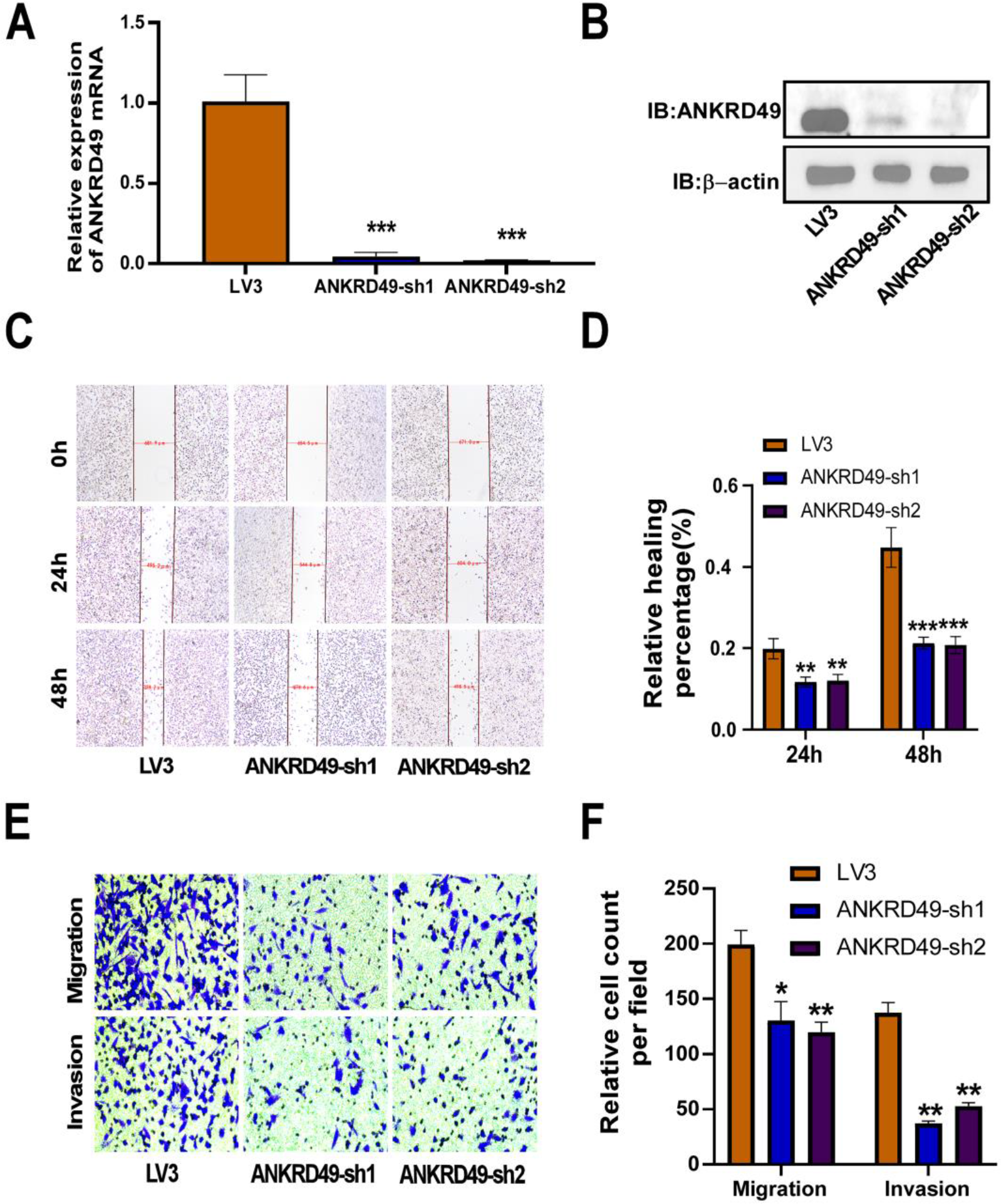
Knockdown of ANKRD49 inhibits migration, invasion of H1703 cells. (A, B) The stable ANKRD49-sh H1703 cells was validated by RT-qPCR and Western blot. β-Tubulin served as an internal control. (C, D) Wound healing assay was performed to detect migration of ANKRD49-sh H1703 cells, representative images were taken at 40× magnification at 0, 24 and 48 h. (E, F) Transwell migration and invasion assays were conducted to evaluate the migration and invasion of ANKRD49-sh H1703 cells. Representative images were taken at 200× magnification. Data are expressed as means ± standard deviation. \**P*<0.05, \*\**P*<0.01, \*\*\**P*<0.001 vs LV3 group.

**Fig. 7.**
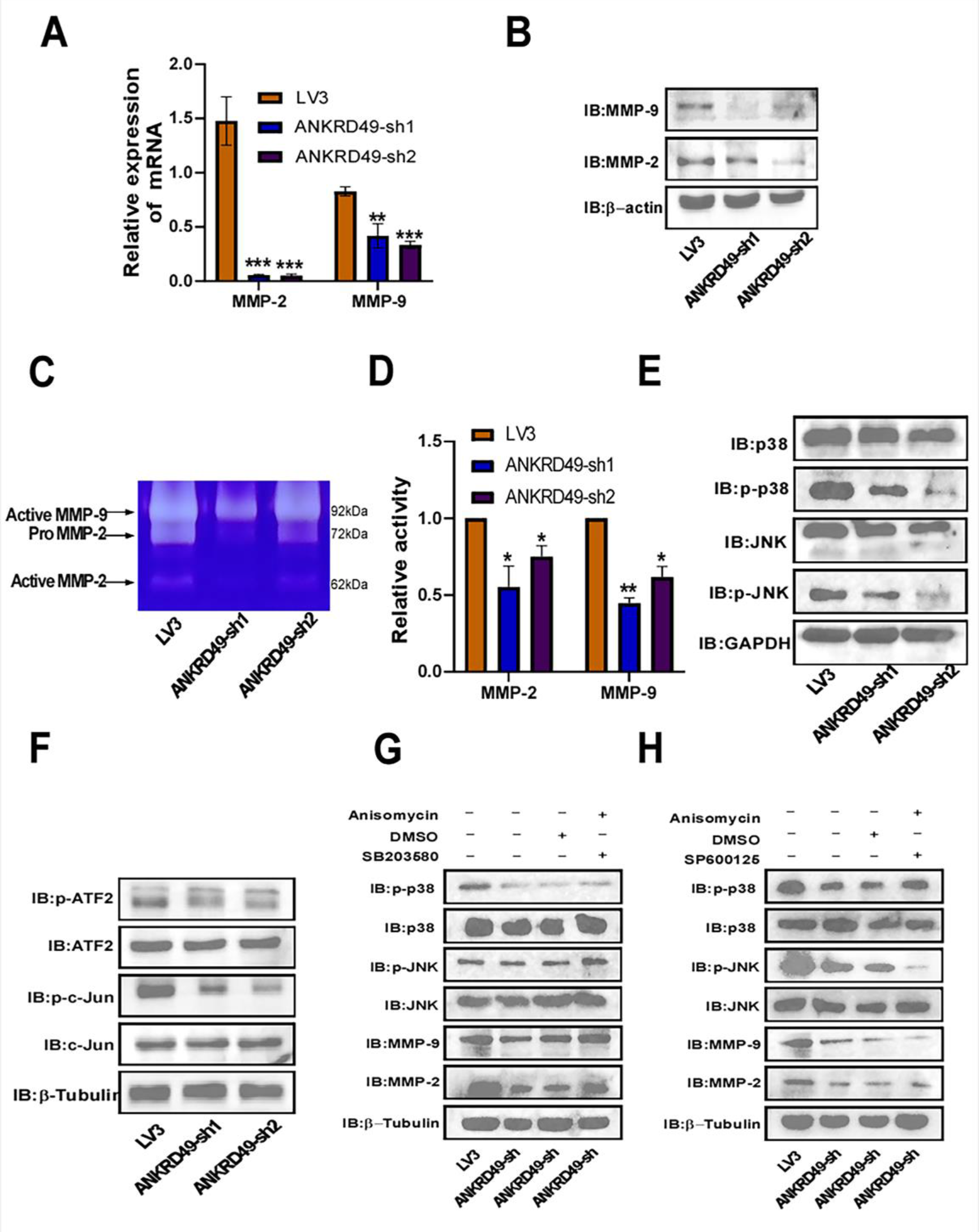
Knockdown of ANKRD49 downregulates MMP-2/MMP-9 expression of H1703 cells. (A, B) The mRNA and protein levels of MMP-2 and MMP-9 in ANKRD49-sh H1703 cells were analyzed by RT-qPCR and Western blot. (C, D) The activities of MMP-2 and MMP-9 in ANKRD49-sh H1703 cells were detected by gelatin zymography. (E) The MAPK proteins in ANKRD49-sh H1703 cells were detected by Western blot. GAPDH served as an internal control. (F) The levels of p-ATF2 and p-c-Jun in ANKRD49-sh H1703 cells were measured by Western blot. (G, H) The levels of JNK, p-JNK, p38, p- p38, MMP-2 and MMP-9 in ANKRD49-sh H1703 cells treated with SP600125, SB203580, Anisomycin or DMSO were measured by Western blot. β-Tubulin served as an internal control. Data are expressed as means ± standard deviation. \**P*<0.05, \*\**P*<0.01, \*\*\**P*<0.001 vs LV3 group.

### 2.6 ANKRD49 potentiates migration and invasion of H1299 and H1703 cells in nude mice

To further explore the effect of ANKRD49 on the migration of NSCLC cells, ANKRD49-OE or LV5 H1299 cells and ANKRD49-sh or LV3 H1703 cells were injected intravenously into the tail of nude mice and maintained for 4 weeks. The results appeared that ANKRD49-OE enhanced the incidence of lung metastasis and the number of metastatic foci (Fig. 8A, B). Downregulation of ANKRD49 substantially reduced the incidence of lung metastasis and number of metastatic foci (Supplementary Fig. S3A, B). Images of the mouse lungs are shown in Supplementary Fig. S4. HE staining also illustrated that more tumor cells appeared in perivascular and peribronchial regions in lung tissues from mice injected with ANKRD49-OE H1299 cells than in control mice (Fig. 8C). Nevertheless, the distribution of tumor cells in the lung tissues of mice injected with ANKRD49-sh or LV3 showed the opposite trend (Supplementary Fig. S3C).

**Fig. 8.**
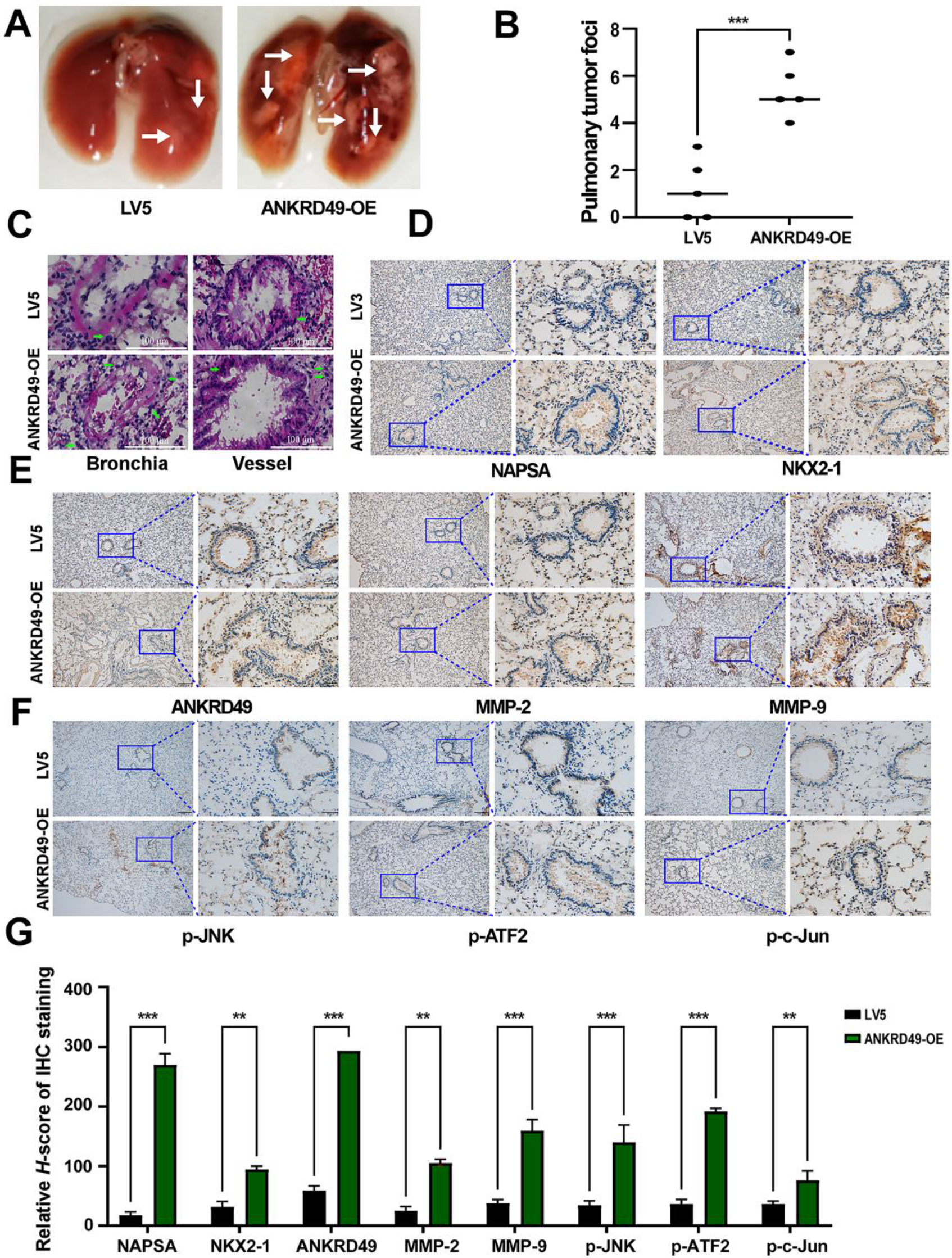
Overexpression of ANKRD49 promotes the migration and invasion of H1299 cells in nude mice. (A) Representative lung images of mice injected with ANKRD49-OE-H1299 or LV5-H1299 cells. White arrows manifest the metastasis nodules on the lung. (B) Statistical analysis of the number of metastasis nodules on the lung was illustrated. (C) Representative images of HE staining for lung metastases. Greenarrows indicated metastatic H1299 cells. (D) Representative images of IHC staining for NAPSA and NKX2-1. Scale bars represent 100 μm. (E, F) Representative images of IHC staining for ANKRD49, MMP-2, MMP-9, p-JNK, p-ATF2 or p-c-Jun. Scale bars represent 100 μm. (G) Quantitative analysis of IHC staining. \*\**P*<0.01, \*\*\**P*<0.001 vs LV5 group.

NAPSA and NKX2-1 are markers for LUAD(Jones et al., 2018; Wu et al., 2020; Yang et al., 2012), whereas p63 is a marker for LUSC(Nobre et al., 2013). Immunohistochemical staining illustrated that the distribution of ANKRD49-positive cells as well as NAPSA-positive, NKX2-1-positive, or p63- positive cells was consistent with the results of HE staining (Fig. 8D; Supplementary Fig. S3D). In addition, immunohistochemical staining showed that the levels of ANKRD49, MMP-2, MMP-9, p-JNK, p-c-Jun, and p-ATF2 were elevated in lung tissues from mice injected with ANKRD49-OE H1299 cells (Fig. 8E-G), whereas their levels were decreased in lung tissues of mice injected with ANKRD49-sh H1703 cells (Supplementary Fig. S3E-G), compared to control mice. Correlation analysis displayed that the metastatic rates of the tumor cells were positively correlated with the expression of ANKRD49, MMP-2, MMP-9, p-JNK, p-c-Jun or p-ATF2 (Supplementary Fig. S5). In addition, MMP-2, MMP-9, p-JNK, p-ATF2, and p-c-Jun levels were positively correlated with the levels of ANKRD49 (Supplementary Fig. S6). In summary, the *in vivo* assay demonstrated that ANKRD49 promoted the invasion and migration of H1299 and H1703 cells by upregulating MMP-2/MMP-9 expression via activating of the JNK- ATF2/c-Jun pathways.

## 3. Discussion

The ANKRD49 protein contains four ankyrin repeat motifs involved in multiple protein-protein interactions in diverse life activities(Wang et al., 2015). It has been documented that ANKRD49 is highly expressed in several carcinomas, including gastric cancer and malignant gliomas. In human malignant gliomas, ANKRD49 reduces cellular apoptosis and facilitates cell cycle progression to promote the proliferation of glioma cells(Hao et al., 2017). It also has been documented that ANKRD49 may serve as an independent prognostic indicator of gastric cancer since it’s high expression(Liu et al., 2018). In a conjoint analysis of gene expression omnibus, computational NCI-60 microarray databases, and an RT-qPCR assay based on sixty-nine patients, *ANKRD49* was identified as an invasion-correlated gene in NSCLC(Hsu et al., 2009). Our previous study using tissue microarray analysis of 160 NSCLC patients which contains 80 LUAD cases and 80 LUSC cases combined with IHC staining showed that ANKRD49 is highly expressed in cancerous lung tissues and correlates with lymph node metastasis, distal metastasis, TNM stage, and differentiation in NSCLC patients. Furthermore, higher ANKRD49 expression was associated with poor OS in patients with NSCLC(Li et al., 2022; Liu et al., 2022). These data implied that ANKRD49 participates in the occurrence and development of NSCLC.

Previously, we used A549 cells, a LUAD cell line, to explore the function of ANKRD49 and found that ANKRD49 promoted the metastasis of A549 cells via upregulation of MMP- 2 and MMP- 9 in a p38/ATF-2 pathway- dependent manner (Liu et al., 2022). In consideration of different subtypes of NSCLC as well as heterogeneity of tumor cells, the function of ANKRD49 in NSCLC progression need further clarification. Herein, we used a lentivirus-mediated overexpression vector to upregulate ANKRD49 expression in H1299 cells, another LUAD cell line, and found that ANKRD49 potentiated the invasion and metastasis of H1299 cells. Furthermore, we utilized lentivirus-mediated shRNA, a loss-of-function strategy, to downregulate ANKRD49 expression in H1299 and H1703 cells (a LUSC cell line), and discovered that knockdown of ANKRD49 suppressed the invasion and migration of these cells. The migration and invasion of tumor cells into surrounding healthy tissues and vasculature are the crucial initial step in tumor metastasis, enabling the tumor to colonize distant organs and spread the disease(Duff and Long, 2017). It is widely acknowledged that metastasis is a crucial cause of cancer-related death, and cell migration is a sophisticated, multistep process(Polacheck et al., 2013). During this multistep process, degradation of the extracellular matrix (ECM) is an early event in tumor metastasis(Mikami et al., 2014). MMPs belong to a family of proteolytic enzymes with many physiological roles, including ECM modification, which accelerates cell migration(Wells et al., 2015). Among the MMP family, MMP- 2 and MMP-9 are documented as substrate-specific gelatinases that are pivotal in ECM degradation(Yang et al., 2019) and are advantageous for invasion and metastasis of tumor cells, leading to tumorigenesis(Jacob et al., 2013). Our study demonstrated that ANKRD49 promotes the expression and activity of MMP-2 and MMP-9, which enhances the migration and invasion of NSCLC cells.

The JNK and p38 MAPK signaling pathways are characterized by kinase cascades in cancerous biology and play a central role in the carcinogenesis and maintenance of cancers(Aggarwal et al., 2019; Wagner and Nebreda, 2009). JNK and p38 MAPK have been identified as crucial mediators of migration and invasion in various tumors(Ou et al., 2021; Wang et al., 2019). In this study, the phosphorylation of JNK and p38 was elevated in ANKRD49 ectopic expressed H1299 cells and decreased in ANKRD49 downregulated H1299 and H1703 cells. So, we speculated that the JNK and p38 MAPK pathways were involved in ANKRD49-induced cellular migration and invasion. However, unlike the JNK inhibitor, the p38 MAPK inhibitor did not induce a decline in the levels of MMP-2 and MMP-9. These data demonstrate that the JNK signaling pathway participates in ANKRD49-induced migration and invasion of H1299 and H1703 cells. Activated p38 signaling pathway maybe involved in other processes such as tumor immunity(Lin et al., 2022) rather than metastasis in current settings. The present results were inconsistent with those in A549 cells, a possible reason was that these three types of cell lines have diverse histology and genetic backgrounds. A549 cells have been tested for the KRAS and KEAP1 (kelch like ECH associated protein 1) missense mutation, STK11 (serine/threonine kinase 11) nonsense mutation, wild type p53 expression, while H1299 cells have a homozygous partial deletion of the p53 protein, and lack expression of p53 protein(Hinz et al., 2021). Of course, how these different genetic backgrounds affect p38 signaling pathway remains to be further studied.

Subsequently, we sought to understand how JNK regulates MMP-2 and MMP-9 expression. MMP-2 or MMP-9 transcription is reportedly regulated by various transcription factors, including AP-1, SP-1, NF-κB, and CREB(Li et al., 2020b). ATF2 and c-Jun are components of the dimeric transcription factor AP-1 and play crucial regulatory roles in many biological processes(Lopez-Bergami et al., 2010). Both ATF2 and c-Jun can be phosphorylated by JNK and subsequently translocated into the nucleus to function as transcription factors(Pant et al., 2016). The phosphorylation of ATF2 and c-Jun, as well as their nuclear distribution, were elevated in ANKRD49 overexpressed H1299 cells, and downregulation of ANKRD49 inhibited this tendency. It has been reported that in the absence of c-Jun, the monomers of ATF2 continuously shuttle between the nucleus and cytoplasm because it contains a nuclear export signal (NES) and two nuclear localization sequences (NLS), while the homodimers of ATF2 are predominantly distributed in the cytoplasm(Watson et al., 2017). To function as a transcription factor, ATF2 needs to form a heterodimer with c-Jun, which masks NES and prevents ATF2 nuclear export(Lindaman et al., 2013). Herein, we found that ANKRD49 boosted the interaction between ATF2 and c-Jun in the nucleus through Co-IP and immunofluorescence staining assays. It is well established that the promoter of MMP- 2/MMP-9 contains the binding site of c-Jun(Tombulturk et al., 2019), yet the binding sites of ATF2 in the promoter region of MMP-2 and MMP-9 have not been revealed. In the present study, a CHIP assay was carried out, and it was discovered that ATF2 binding sites exist in -1795∼-1788 of the MMP-2 promoter region and -2042∼-2395 of the MMP-9 promoter region. ANKRD49 facilitated binding of ATF2 to the promoter regions of MMP-2 and MMP-9. In the end, our *in vitro* tests confirmed the ANKRD49’s function which was revealed *in vitro* assays. Consequently, we conclude that ANKRD49 mediates MMP-2/MMP-9 via ATF2/c-Jun heterodimers, which are activated by JNK to promote the migration and invasion of NSCLC cells.

The present study has several unaddressed issues. The role of activated p38 MAPK in ANKRD49- OE H1299 or ANKRD49-sh H1703 cells remains unclear. The mechanism by which ANKRD49 establishes contact with JNK or p38 MAPK remains unknown. Accordingly, more in-depth research is needed to elucidate the function and underlying mechanisms of ANKRD49 in NSCLC progression.

In summary, we identified that ANKRD49 potentiates the migration and invasion of H1299 and H1703 cells by activating the JNK-ATF2/c-Jun signaling cascade to upregulate MMP2 and MMP-9 expression. ANKRD49 may serve as an independent predictor for prognosis evaluation of NSCLC and be a novel and promising cancer therapeutic target. The ANKRD49–JNK-ATF2/c-Jun-MMPs axis promoted NSCLC cells metastasis and ANKRD49 could be a potential therapeutic target for patients with NSCLC.

## 4. Materials and methods

### 4.1 Human tissue samples

Nine pairs of fresh NSCLC tissues and the corresponding adjacent normal tissues were collected from patients who underwent surgery at the First Hospital of Shanxi Medical University from October 2020 to December 2020. No patients in this study received other treatment before surgery.

### 4.2 Cell lines and cells culture

The human bronchial epithelial cell line (HBEC) and NSCLC cell lines H1299, A549, H446, H1460, Calu-3, H1703, sk-MES-1 were obtained from the Cell Culture Center of the Chinese Academy of Medical Sciences (Beijing, China). These NSCLC cell lines were cultured in Advanced RPMI 1640 medium (Seven, Beijing, China) supplemented with 10% fetal bovine serum (Every Green, Zhejiang, China) and incubated at 37°C in 5% CO2.

### 4.3 Establishment of ANKRD49 stably expressed cell lines

H1299 and H1703 cells were infected with lentivirus-5 ANKRD49 or lentivirus-3 ANKRD49-shRNA and corresponding vectors (Gene Pharma, Shanghai, China) according to the manufacturer’s protocols to construct the stable ANKRD49 overexpression (ANKRD49-OE) or ANKRD49 knockdown (ANKRD49-sh) groups. Meanwhile, the controls for overexpression (LV5) and knockdown (LV3) were also established. After infection of lentivirus, cells were incubated for more than 48 h and puromycin (2 μg/ml) was added to screen the stable cells. LV3 and ANKRD49-sh sequences were listed in Supplementary Table 1.

### 4.4 Reagents and antibodies

Puromycin (P816466) was purchased from MACKLIN (Shanghai, China). Anisomycin (GC11559) was purchased from GLPBIO (Shanghai, China). The p38 mitogen-activated protein kinase (MAPK) inhibitor (HY-10256, SB203580) and JNK inhibitor (HY-12041, SP600125) were purchased from MedChemExpress (New Jersey, USA). MMPs inhibitor (SF4180, ilomastat) was purchased from Beyotime Biotechnology (Shanghai, China). Rabbit anti-ANKRD49 primary antibody (Proteintech, Cat# 25034-1-AP, RRID:AB_2879860), rabbit anti-MMP-2 monoclonal antibody (Cat# bs-4599R, RRID:AB_11083963), rabbit anti-MMP-9 monoclonal antibody (Cat# bs-0397R, RRID:AB_10853038), rabbit anti-c-Jun monoclonal antibody (Cat# bs-0670R, RRID:AB_10857880), rabbit anti-p-ATF2 monoclonal antibody (Cat# bs-3033R, RRID:AB_10883830) and Alexa Fluor^®^ 488-conjugated goat anti-rabbit IgG (Cat# bs-0295G-A488, RRID:AB_10893781) were obtained from Bioss Biotechnology (Beijing, China). The mouse anti-p-c-Jun monoclonal antibody (Cat# 558036, RRID: AB_2249448) was obtained from BD Biosciences (New Jersey, USA). Rabbit anti-p38 monoclonal antibody (Cat# 8690, RRID: AB_10999090), rabbit anti-p-p38 monoclonal antibody (Cat# 4511, RRID: AB_2139682), rabbit anti-ERK monoclonal antibody (Cat# 4695, RRID: AB_390779), rabbit anti-p-ERK monoclonal antibody (Cat# 4376, RRID: AB_331772), rabbit anti-SAPK/JNK monoclonal antibody (Cat# 9258, RRID: AB_2141027), and rabbit anti-p-JNK monoclonal antibody (Cat# 4668, RRID: AB_823588) were purchased from Cell Signaling Technology (Danvers, MA, USA). Rabbit anti-ATF2 monoclonal antibodies (Cat# BS1022, RRID: AB_1664115), rabbit anti-β-actin monoclonal antibodies (Cat# AP0060, RRID: AB_2797445), rabbit anti-GAPDH monoclonal antibodies (Cat# AP0063, RRID: AB_2651132), rabbit anti-Histone-H3 monoclonal antibodies (Cat# BS7675), rabbit anti-PARP monoclonal antibodies (Cat# BS7190), and rabbit anti-β-tubulin monoclonal antibodies (Cat# AP0064, RRID: AB_2797447) were acquired from Bioworld (Minnesota, USA). Rabbit anti-ATF2 polyclonal antibodies (Cat# D155200) and rabbit anti-p-ATF2 polyclonal antibodies (Cat# D155010) were purchased from Sangon Biotech (Shanghai, China). Anti-rabbit IgG (H+L) (Cat# BA1050, RRID: AB_2904507) and anti-mouse IgG (H+L) (Cat# BA1054, RRID: AB_2734136) were purchased from Boster Biological Technology (Wuhan, China). Dylight 649 and goat anti-mouse IgG (Cat# 610-143-002, RRID: AB_11182582) were obtained from Rockland (Philadelphia, USA).

### 4.5 Western blot

Total protein from cells or tissues was extracted using SDS lysis buffer containing 2% SDS, 10% glycerol, and 50 mM Tris-HCl (pH 6.8), and quantified using the Enhanced BCA Protein Assay Kit (Beyotime Biotechnology). Proteins (30 μg) were loaded onto a 12% SDS-PAGE gel, separated, and transferred onto polyvinylidene fluoride (PVDF) membranes. The membranes were blocked using 5% skimmed milk, followed by probing with the indicated primary antibodies at 4°C overnight, and then incubated with an anti-rabbit/mouse HRP-conjugated IgG at room temperature for 1 h. Blots were developed using the ECL kit (Seven, Beijing, China) and detected using the ChemiDoc imaging system (Bio-Rad, California, USA).

### 4.6 Real-time polymerase chain reaction

Total RNA from NSCLC cells, nine pairs of human NSCLC tissues, and matched adjacent normal tissues were prepared using the Total RNA Extraction Kit (Promega, Madison, WI, USA) following the manufacturer’s instructions. RT-qPCR was performed using the RT-PCR Kit (Seven, Beijing, China) on the Quant Studio 3 (Thermo Fisher Scientific, Massachusetts, USA), according to the manufacturer’s procedures. All the primers are listed in Supplemental Table 2. β-actin was used as the control, and the relative mRNA expression was defined using the 2 ^-ΔΔCt^ method.

### 4.7 Gelatin zymography

MMP-2 and MMP-9 activities were evaluated using gelatin zymography. 5×10^5^ cells were pretreated with serum-free medium for 48 h. The medium was collected, centrifuged at 1,500× g for 15 min at 4°C, and mixed with non-reducing sample buffer for electrophoresis on a polyacrylamide gel containing 0.1% (w/v) gelatin. Proteolysis was assessed as a white zone in the dark blue fields. The 62 kDa and 92 kDa bands corresponded to active MMP-2 and MMP-9, respectively. The gelatinase activity of MMP-2 and MMP-9 was illustrated via the area of the clear zone in the dark blue gel and analyzed using ImageJ analysis software (National Institute of Health, Maryland, USA).

### 4.8 Cell counting kit-8 (CCK-8) assay

The CCK-8 assay was performed to evaluate cellular proliferation, according to the manufacturer’s protocols. Briefly, ANKRD49-OE or LV5 H1299 cells, ANKRD49-sh, or LV3 H1703 cells were cultured in 96-well plates at a density of 1×10^3^ cells per well in a 5% CO2 incubator at 37°C. Next, 100 μl serum-free medium containing 10 μl CCK-8 reagent (Seven, Beijing, China) was added to each well, followed by another 1 h incubation. The absorbance of the cells was measured at 450 nm using a microplate reader (Molecular Devices, California, USA).

### 4.9 Colony formation assay

ANKRD49-OE or LV5 H1299 cells, ANKRD49-sh, or LV3 H1703 cells were plated in six-well plates at a density of 5×10^2^ cells/well and cultured at 37°C for 1 week. After washing with PBS, the cells were fixed with methanol and stained with 0.1% crystal violet solution. The colonies were photographed and quantified using a light microscope (Eclipse Ts2; Nikon, Tokyo, Japan).

### 4.10 Wound healing assay

Cells (2×10^5^) were seeded into six-well plates and cultured for 24 h. Wounds in the confluent cells were created using a 200 μl pipette tip, followed by washing the cells with PBS to remove any floating cells. Next, complete medium was added and the wound margins were photographed. Subsequently, wound healing was photographed at 24 h and 48 h using a Nikon photomicroscope (Eclipse Ts2, Nikon, Tokyo, Japan). Migratory ability was calculated by measuring the distance between different groups of cells simultaneously.

### 4.11 Migration and invasion assays

Cells (2×10^4^) were seeded into the upper chamber of 24-well inserts (8.0 µm, Corning, California, USA). The upper chamber was pre-coated with Matrigel (BD Biosciences, California, USA), and a medium containing 20% FBS was added to the lower chamber for the invasion assay. After incubation at 37°C for 24 h, the remaining cells in the upper chamber were scraped off and the cells on the lower side of the chamber were fixed, stained, and viewed using a Nikon photomicroscope (Eclipse Ts2). Data were analyzed by counting at least five random fields.

### 4.12 Cytoplasmic and nuclear protein extraction

The subcellular proteins in ANKRD49-OE or LV5 H1299 cells were prepared using a subcellular structure cytoplasm and nucleus extraction kit (Bioss Biotechnology). The protein samples (20 μg) were subjected to Western blot with p-ATF2 or p-c-Jun antibodies.

### 4.13 Immunofluorescence

H1299 cells were transfected with pMSCVpuro-ANKRD49 or the vector for 48 h, and 5×10^4^ cells were seeded onto sterile coverslips in a twelve-well plate for another 24 h. The medium was removed, and cells were fixed with 4% paraformaldehyde and then permeabilized with 0.01% Triton X-100 at room temperature for 10 min. Cells were incubated with anti-p-ATF2 antibody or anti-p-c-Jun antibody at 4°C overnight, and then probed with Alexa Fluor^®^ 488-conjugated goat anti-rabbit IgG or Dylight 649 goat anti-mouse IgG at room temperature for 1 h in darkness. Nuclei were stained with DAPI (Beyotime) for 5 min, and fluorescence signals were observed using a fluorescence microscope (Eclipse Ts2).

### 4.14 Co-Immunoprecipitation (Co-IP)

Nucleus proteins of the ANKRD49-OE or LV5 H1299 cells (1×10^6^) were extracted and 25 µg of them were precleared using Protein A/G Agarose (Santa Cruz, California, USA). Subsequently, 2 μl rabbit anti-ATF2 or normal rabbit IgG was added to the lysates and incubated at 4°C for 12 h. Next, 20 μl of Protein A/G PLUS Agarose was mixed into the lysates and incubated at 4°C overnight. Agarose was washed three times using wash buffer, and the bound proteins were separated by SDS-PAGE.

### 4.15 Immunohistochemistry (IHC)

Five micrometer-thick sections of formalin-fixed paraffin-embedded lung tissues from nude mice were prepared for IHC staining. Slides were deparaffinized, rehydrated, and microwaved for antigen retrieval in 0.01 M, PH 6.0 sodium citrate buffer. Endogenous peroxidase activity was blocked using 3% H2O2.

Subsequently, sections were incubated with the corresponding primary antibody (1:100) for 45 min at room temperature. Diaminobenzidine (DAB) (Boster, Wuhan, China) was used to visualize peroxidase activity. The immunostaining intensity of ANKRD49, NAPSA, NKX2-1, MMP-2, MMP-9, p63, p-JNK, p-c-Jun, and p-ATF2 was calculated according to the published *H*-score method: *H*-score value = (unstained tumor cells) % × 0 + (weakly stained tumor cells) % × 1 + (moderately stained tumor cells) % × 2 + (strongly stained tumor cells) % × 3.

### 4.16 Chromatin immunoprecipitation assay

The chromatin immunoprecipitation (CHIP) assay was conducted in accordance with the manufacturer’s protocols (Beyotime Biotechnology). The binding site of ATF2/c-Jun in the MMP-2 or MMP-9 promoter region and the position of the CHIP primers are illustrated in Supplementary Fig. S7. Chromatin solutions were sonicated and incubated with anti-p-ATF2, anti-p-c-Jun, or control IgG at 4°C for 12 h. DNA-protein cross-links were reversed, and chromatin DNA was subsequently purified and subjected to PCR analysis. Primers for amplifying the MMP-2 or MMP-9 promoter, which contains the binding site of ATF2/c-Jun, are listed in Supplementary Table 3. The immunoprecipitated DNA was analyzed by qPCR. The qPCR products were resolved on a 1.2% agarose gel and visualized by nucleic acid dye staining.

### 4.17 Animal studies

Ten female BALB/c nude mice and ten male BALB/c nude mice (5~6-week-old) were obtained from the Experimental Animal Center of Shanxi Medical University. The mice were housed under SPF conditions with a 12 h light-dark cycle and ad libitum access to food and water at 23°C with 60% humidity. Female mice were randomly allocated to either LV5 (n=5) or ANKRD49-OE H1299 cells (n=5).

Male mice were randomly allocated to either LV3 (n=5) or ANKRD49-sh H1703 cells (n=5). One week of acclimation, 4×10^6^ cells in 200 μl saline were injected into the lateral tail veins of the nude mice. After the injection, the health of the mice was monitored daily. After 4 weeks, the mice were sacrificed, and whole lungs were collected to count the metastatic nodules on the surface. Lung tissues were fixed in 10% paraformaldehyde for further hematoxylin-eosin (H&E) staining and IHC analysis. All animal experiments were approved by the Animal Ethics Committee of the Shanxi Medical University (approval number SYDL-2020016) and was performed according to the procedures that complied with the ARRIVE guidelines.

### 4.18 Statistical analysis

Correlations between the level of ANKRD49 and the levels of MMP-2, MMP-9, p-JNK, p-ATF2, and p-c-Jun were analyzed using Pearson’s correlation coefficient. A strong correlation was defined as an r-value of ≥ 0.5. SPSS18.0 (IBM, Inc) and GraphPad Prism7.0 (GraphPad Software Inc.) software were used to analyze the data. The analyses were repeated in triplicate. Quantitative data are expressed as the mean ± standard deviation (SD) and analyzed using a Student’s t-test or one-way analysis of variance followed by Tukey’s post hoc test using GraphPad Prism software. A two-sided P-value <0.05 was considered to indicate a statistically significant difference.

## Abbreviations

NSCLC, non-small cell lung cancer; LUAD, lung adenocarcinoma; LUSC, lung squamous cell carcinoma; ANKRD49, ankyrin repeat domain 49; shRNA, short hairpin RNA; MMP, matrix metalloproteinase; ECM, extracellular matrix; MAPK, mitogen-activated protein kinase; ERK, signal-regulated kinases; JNK, c-Jun N-terminal kinase; ATF2, activating transcription factor 2; CO-IP, co-immunoprecipitation; CHIP, chromatin immunoprecipitation; LV, lentivirus; CCK-8, cell counting kit-8; DNA, deoxyribonucleic acid; PCR, polymerase chain reaction; SYBR, synergy brands; GAPDH, glyceraldehyde-3-phosphate dehydrogenase; IHC, immunohistochemistry; RT-qRCR, Real-time quantitative PCR; BCA, bicinchoninic acid; SDS-PAGE, sodium dodecyl sulfate polyacrylamide gel electrophoresis; PVDF, poly vinylidene fluoride membrane; NES, nuclear export signal; NLS, nuclear localization sequence; DAPI, diamidino-phenyl-indole; SPF, Specific Pathogen Free; KEAP1, kelch like ECH associated protein 1; STK11, serine/threonine kinase 11 .

## Acknowledgments

We thank Home for Researchers (www.home-for-researchers.com) for correcting the grammar and spelling.

## Author Contributions

MP and HW conceived and designed the project. JS, JH, CL completed the experiments and analyzed the data. YL collected the human samples. MG, RF and WW analyzed the data from animal experiments. JS and JH wrote the manuscript. HW provided advice and critical comments. The MP was responsible for research supervision and funding acquisition. All authors have read and approved the final manuscript.

## Availability of data and materials

All materials are available by the corresponding author.

## Declarations

### Ethics approval and consent to participate

All tissue samples were obtained in accordance with the Declaration of Helsinki (1975) and were approved by the ethics committee of the First Hospital of Shanxi Medical University (approval number 2019K042). Informed consent was obtained from all participants. All animal experiments were approved by the Animal Ethics Committee of the Shanxi Medical University (approval number SYDL-2020016).

### Consent for publication

All authors consent to publish the paper in Journal of Cell Science

## Supplementary figure legends

**Fig. S1.**
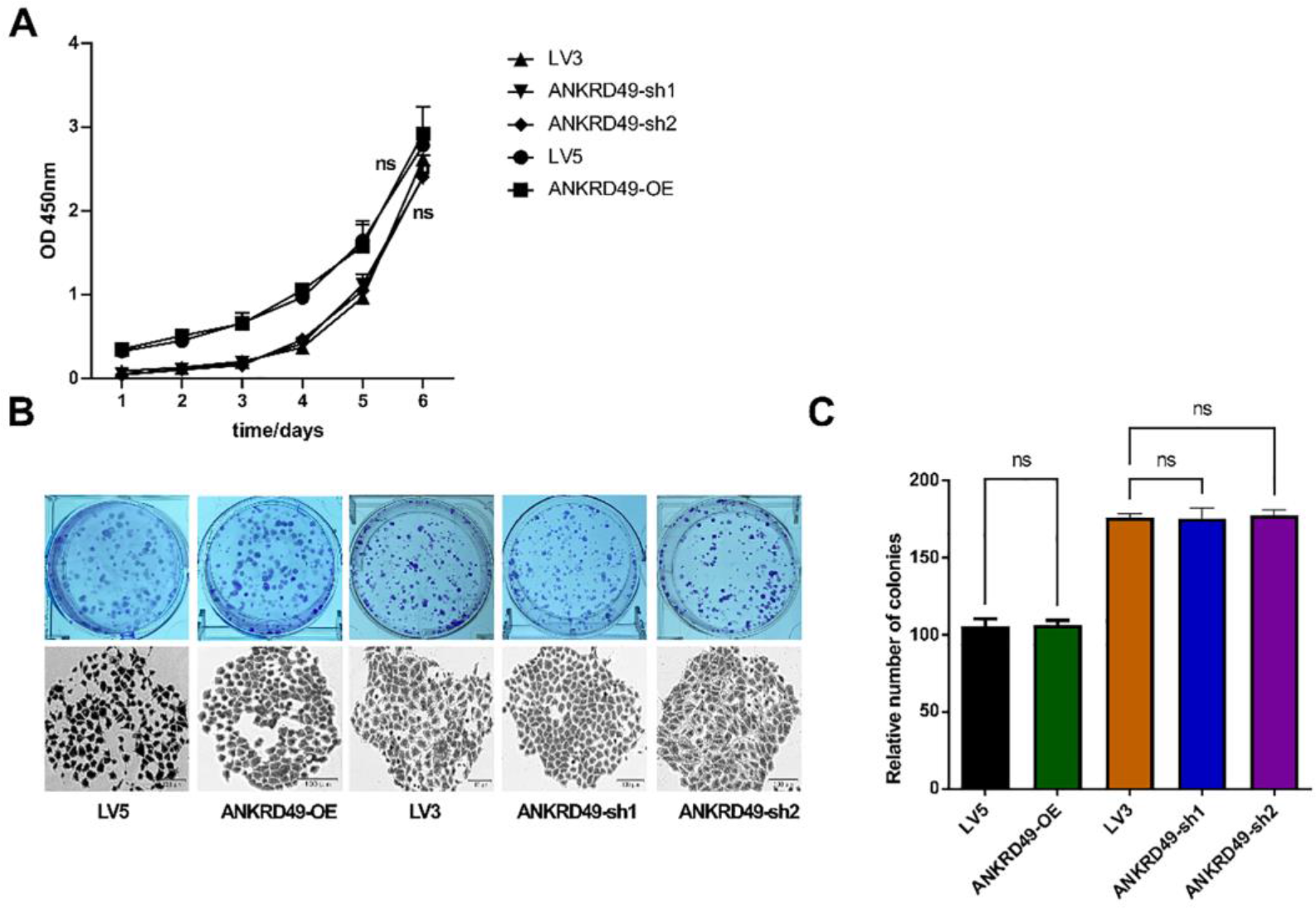
ANKRD49 has no effect on cellular proliferation of H1299 cells. Proliferation assays were performed using CCK8 (A) and colony formation (B, C) in ANKRD49-OE and ANKRD49-sh H1299 cells. All experiments were repeated independently three times. Data are presented as means ± standard deviation. ns, not significant.

**Fig. S2.**
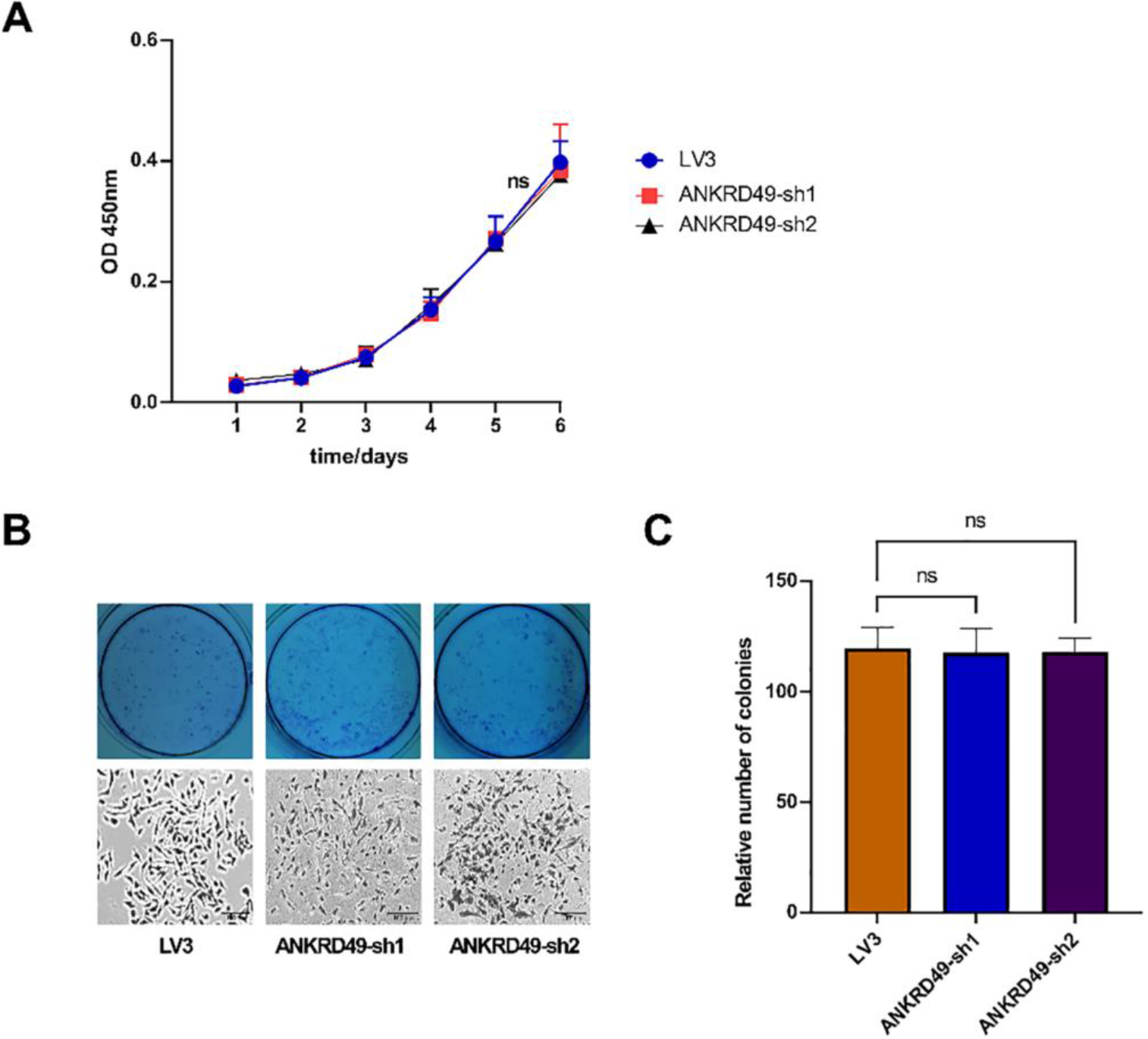
ANKRD49 has no effect on cellular proliferation of H1703 cells. Proliferation assays were conducted using CCK8 (A) and colony formation (B, C) in ANKRD49-sh H1703 cells. All experiments were repeated independently three times. Data are presented as means ± standard deviation. ns, not significant.

**Fig. S3.**
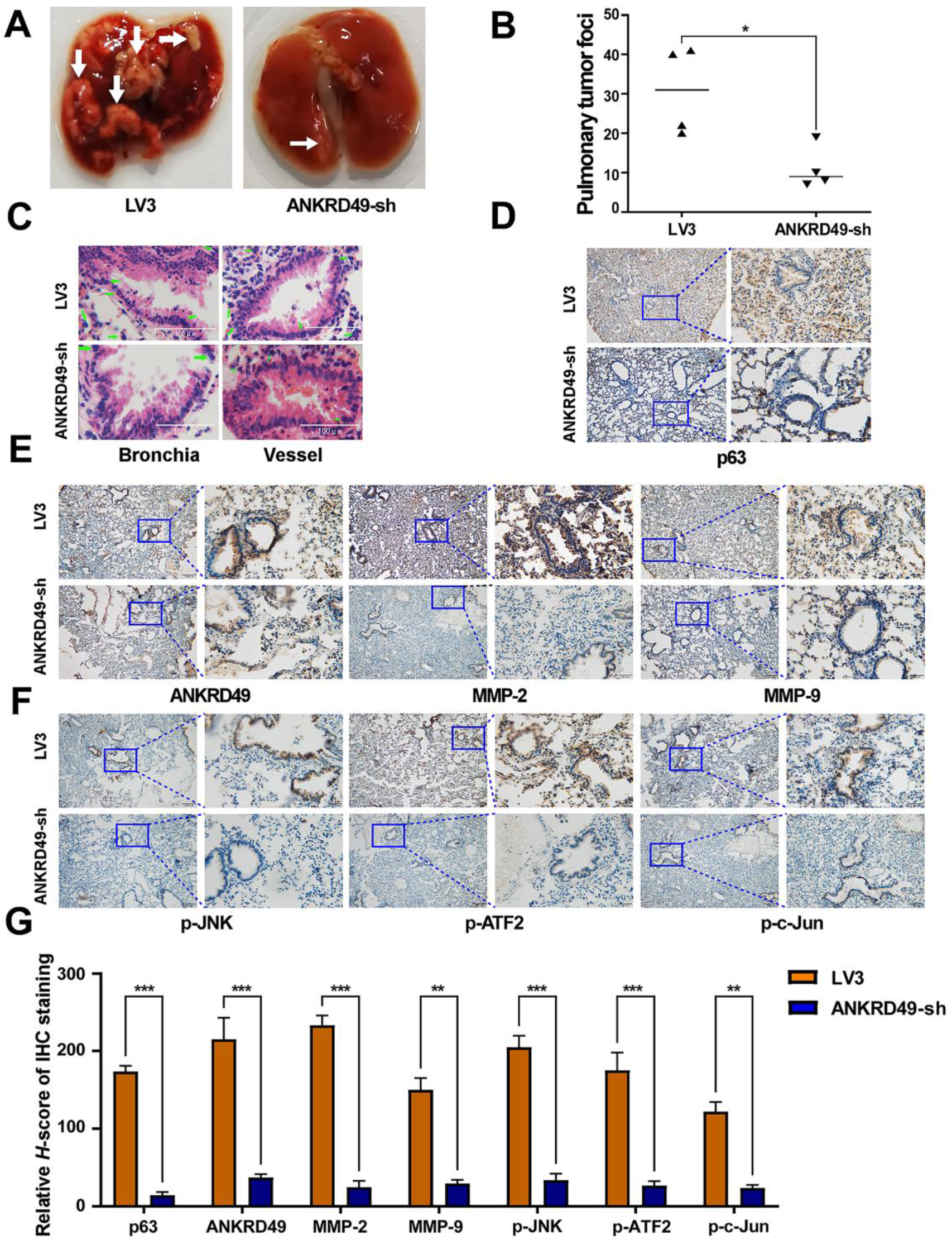
Knockdown of ANKRD49 declines the migration and invasion of H1703 cells in nude mice. (A) Representative lung images of mice injected with ANKRD49-sh or LV3 H1703 cells. White arrows manifest the metastasis nodules on the lung. (B) Statistical analysis of the number of metastasis nodules on the lung was illustrated. (C) Representative images of HE staining for lung metastases. Green arrows indicated metastatic H1703 cells. (D) Representative images of IHC staining for p63. Scale bars represent 100 μm. (E, F) Representative images of IHC staining for ANKRD49, MMP-2, MMP-9, p-JNK, p-ATF2 or p-c-Jun. Scale bars represent 100 μm. (G) Quantitative analysis of IHC staining. \*\**P*<0.01, \*\*\**P*<0.001 vs LV3 group.

**Fig. S4.**
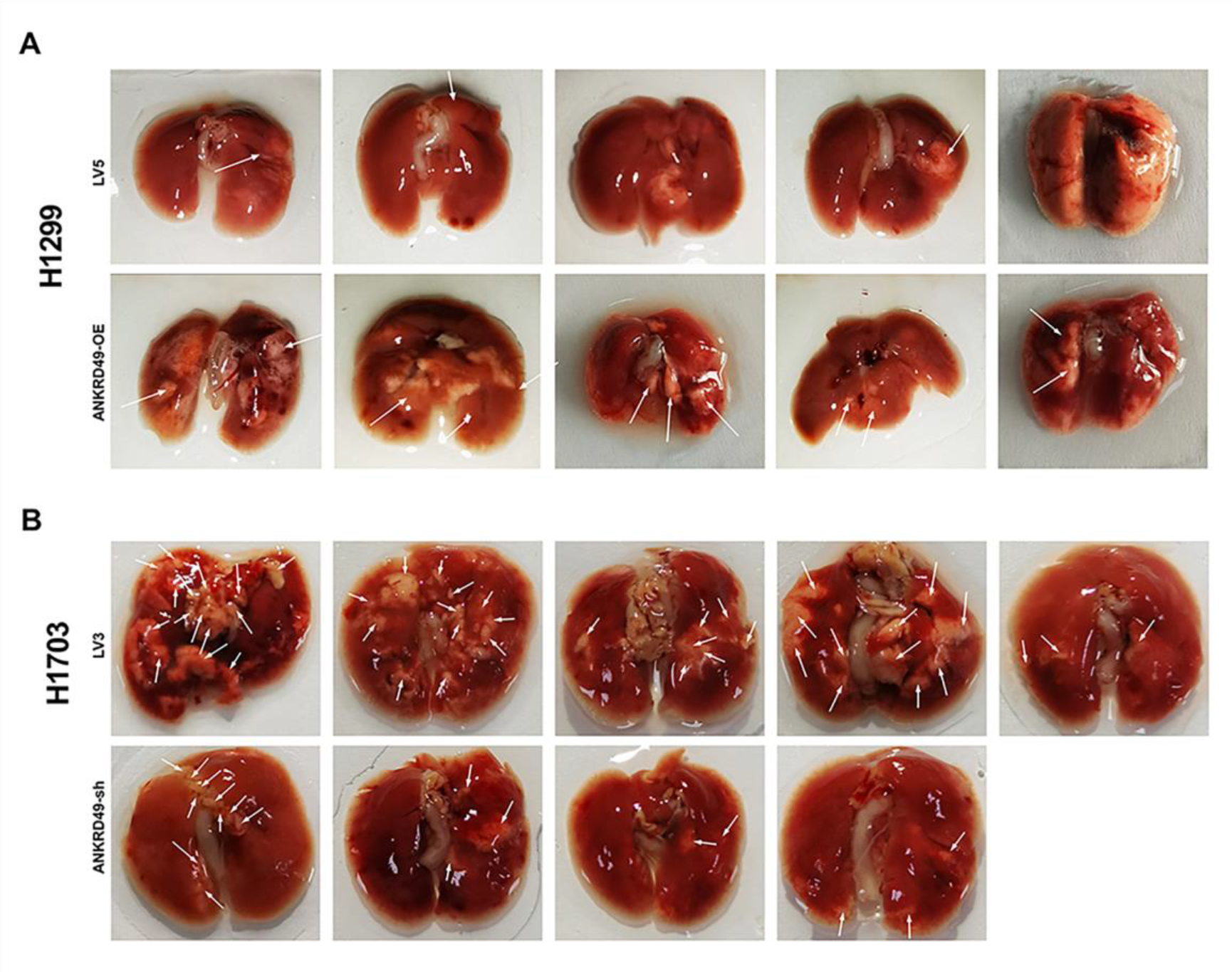
ANKRD49 promotes the lung metastasis of NSCLC cells in nude mice. (A) Images of the lung metastasis nodules in mice injected with ANKRD49-OE or LV5 H1299 cells. (B) Images of the lung metastasis nodules in mice injected with ANKRD49-sh or LV3 H1703 cells. White arrows indicate the metastasis nodules on the lung.

**Fig. S5.**
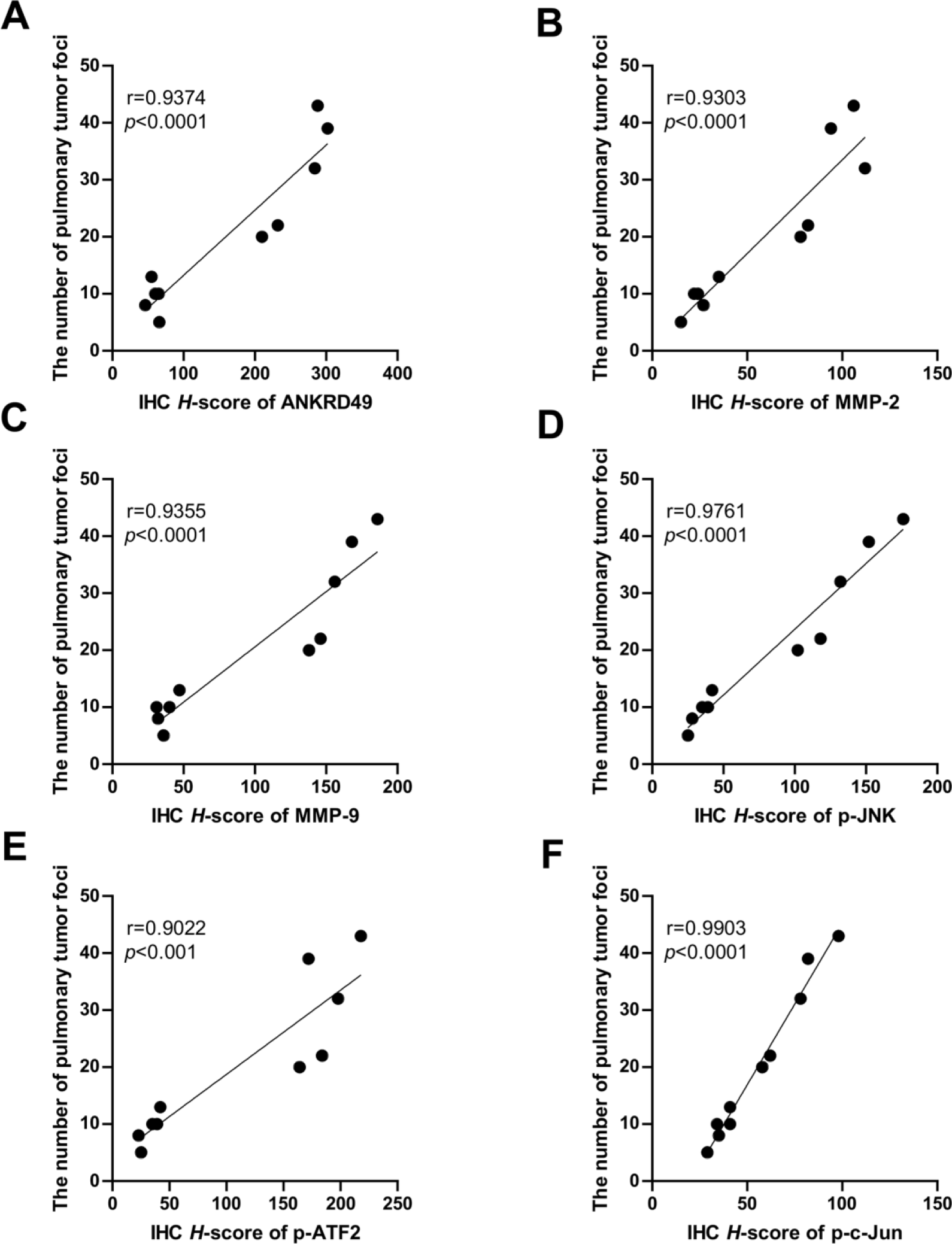
Correlation analysis between H-score of ANKRD49, MMP-2, MMP-9, p-JNK, p-ATF2 or p-c-Jun and lung metastasis in mice injected with ANKRD49-OE H1299 cells. (A) Correlation analysis between H-score of ANKRD49 and lung metastasis in mice injected with ANKRD49-OE H1299 cells. (B) Correlation analysis between H-score of MMP-2 and lung metastasis in mice injected with ANKRD49-OE H1299 cells. (C) Correlation analysis between H-score of MMP-9 and lung metastasis in mice injected with ANKRD49-OE H1299 cells. (D) Correlation analysis between H-score of p-JNK and lung metastasis in mice injected with ANKRD49-OE H1299 cells. (E) Correlation analysis between H-score of p-ATF2 and lung metastasis in mice injected with ANKRD49-OE H1299 cells. (F) Correlation analysis between H-score of p-c-Jun and lung metastasis in mice injected with ANKRD49-OE H1299 cells.

**Fig. S6.**
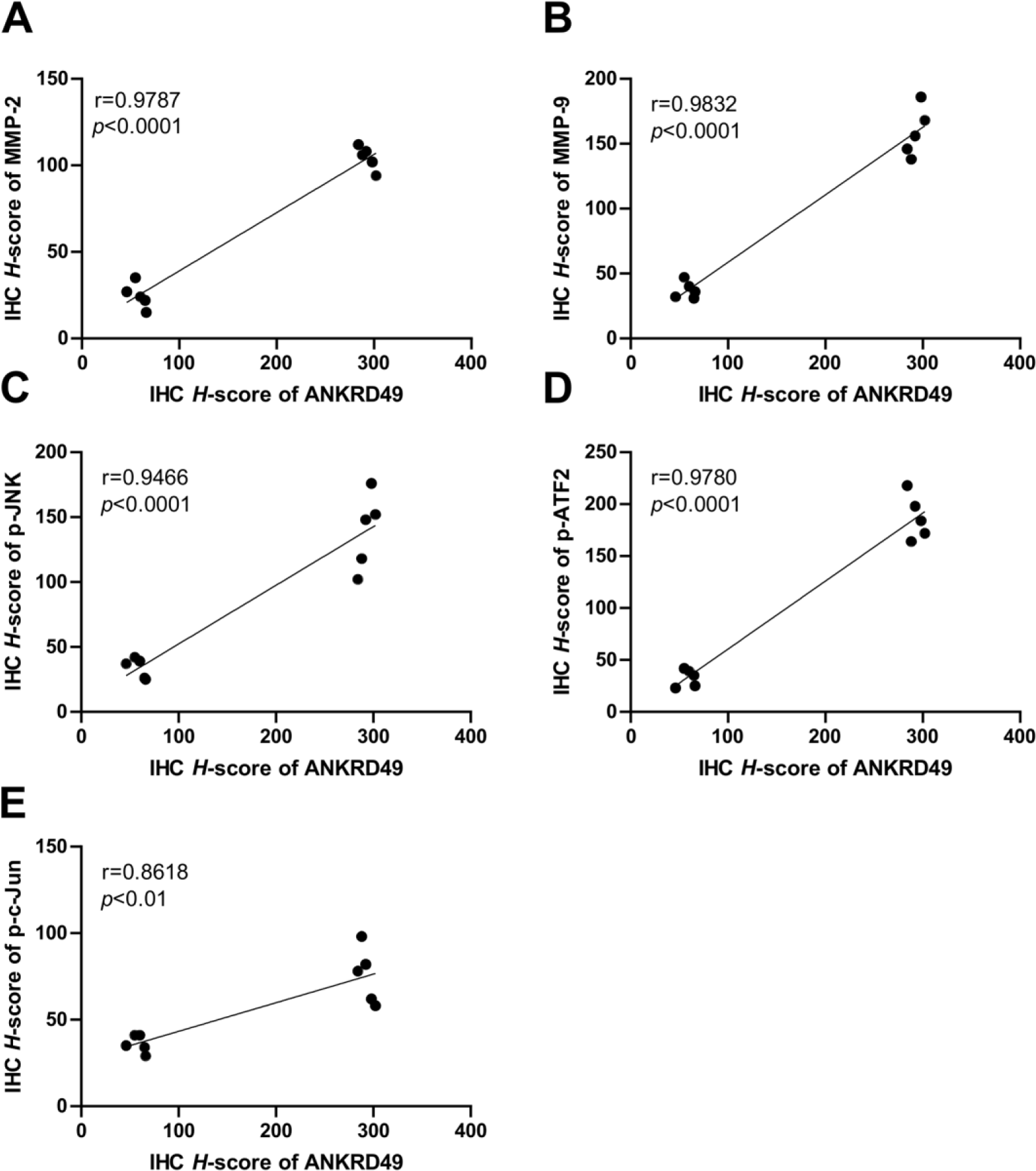
Correlation analysis between H-score of MMP-2, MMP-9, p-JNK, p-ATF2 or p-c-Jun and ANKRD49 in lung tissues of mice injected with ANKRD49-OE H1299 cells. (A) Correlation analysis between H-score of MMP-2 and ANKRD49 in lung tissues of mice injected with ANKRD49-OE H1299 cells. (B) Correlation analysis between H-score of MMP-9 and ANKRD49 in lung tissues of mice injected with ANKRD49-OE H1299 cells. (C) Correlation analysis between H- score of p-JNK and ANKRD49 in lung tissues of mice injected with ANKRD49-OE H1299 cells. (D) Correlation analysis between H-score of p-ATF2 and ANKRD49 in lung tissues of mice injected with ANKRD49-OE H1299 cells. (E) Correlation analysis between H-score of p-c-Jun and ANKRD49 in lung tissues of mice injected with ANKRD49-OE H1299 cells.

**Fig. S7.**
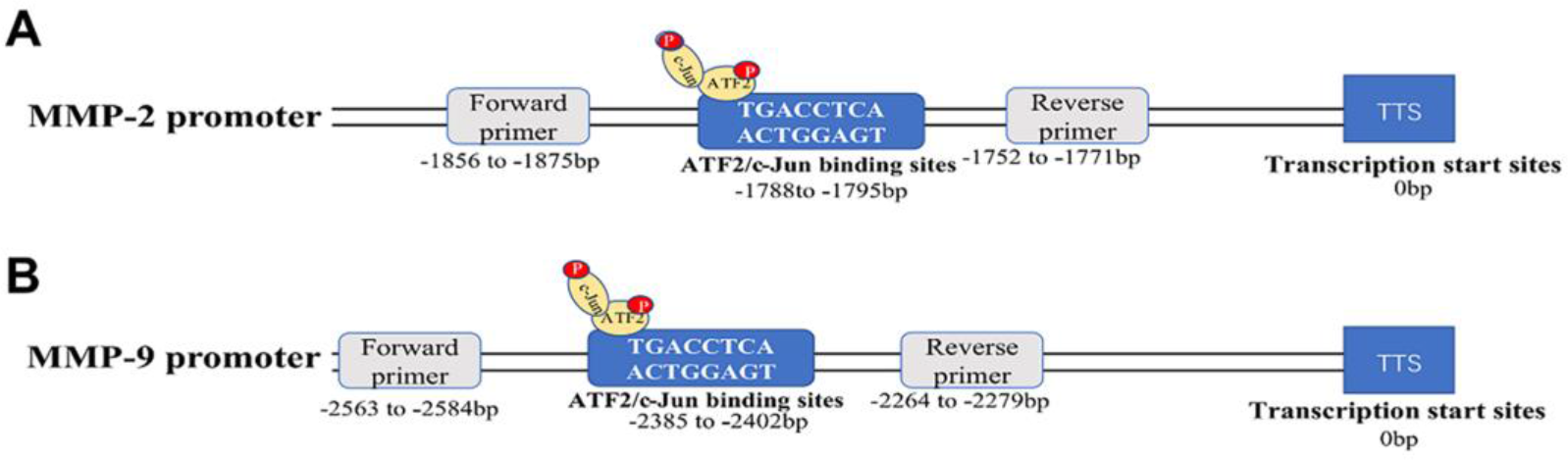
Illustration the binding site of ATF2/c-Jun on MMP-2 or MMP-9 promoter region. (A) illustration the binding site of ATF2/c-Jun on MMP-2 promoter region. (B) illustration the binding site of ATF2/c-Jun on MMP-9 promoter region.

## Supplementary tables

**Supplemental table 1.**
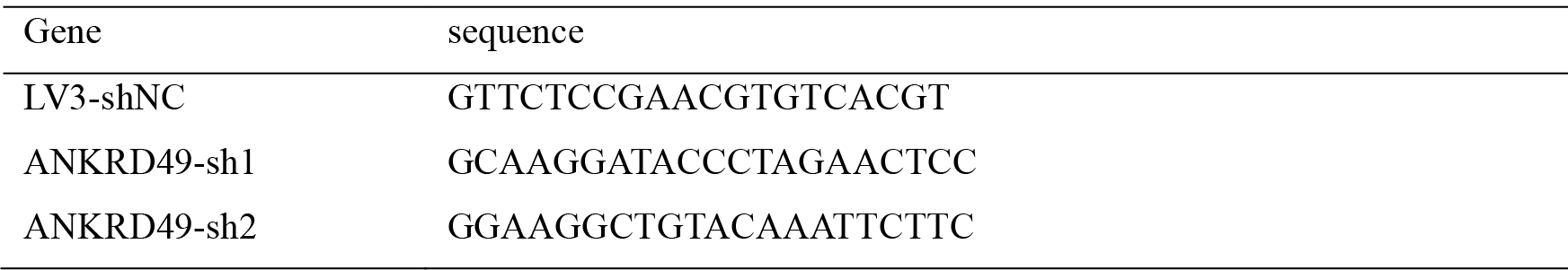
Target sequences for knockdown of ANKRD49.

**Supplemental table 2.**
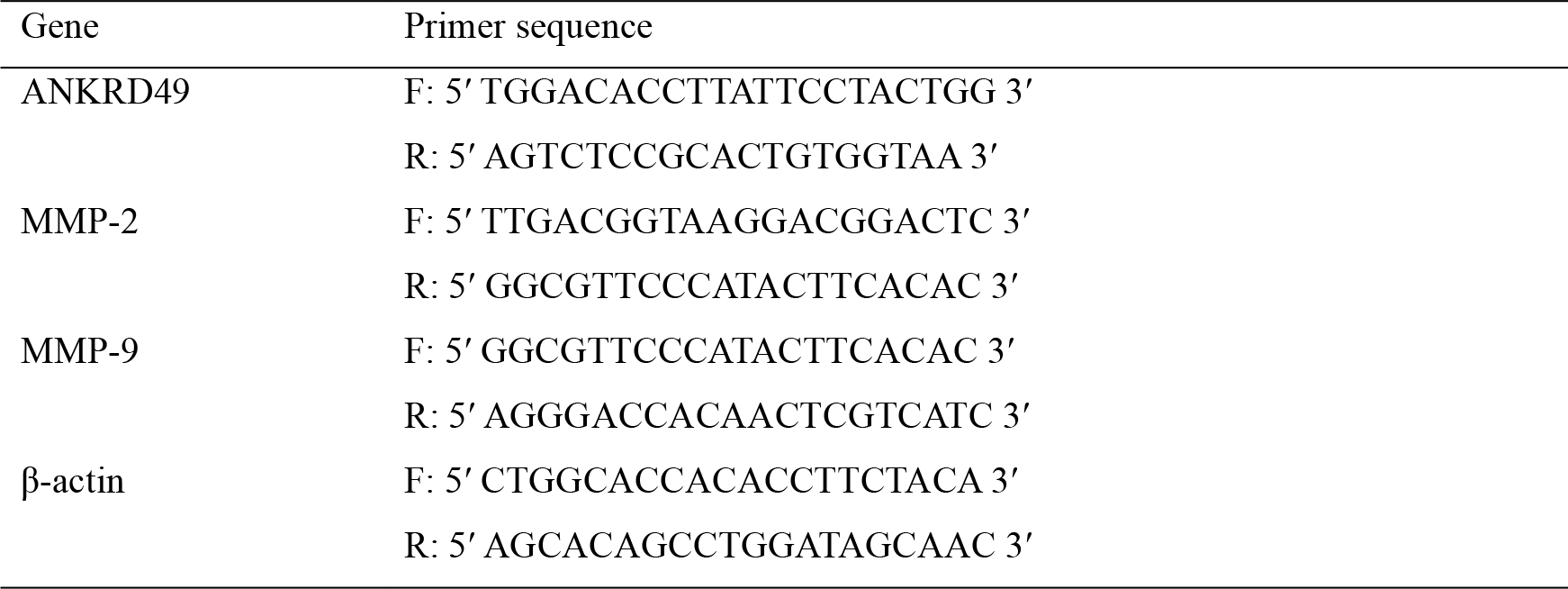
Primers used for real time qPCR in gene expression analysis.

**Supplemental table 3.**
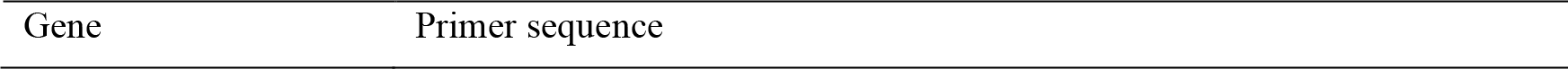
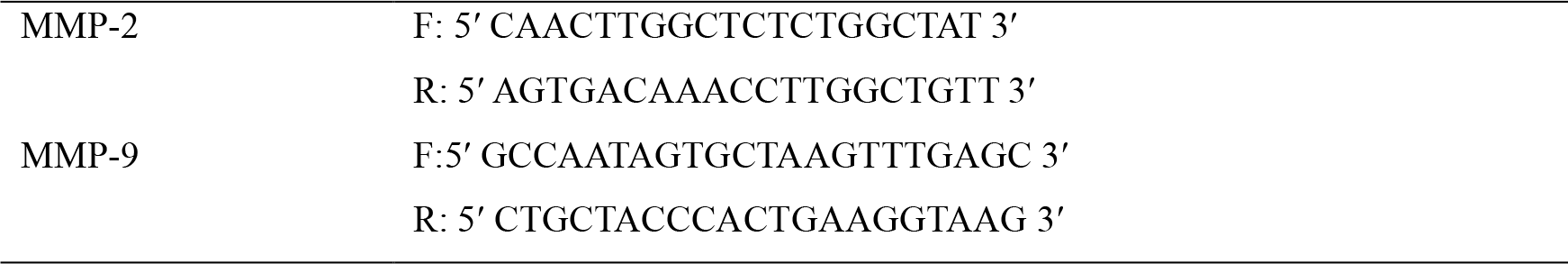
Primers used for ChIP assays.

